# Evolutionary and Comparative analysis of bacterial Non-Homologous End Joining Repair

**DOI:** 10.1101/869602

**Authors:** Mohak Sharda, Anjana Badrinarayanan, Aswin Sai Narain Seshasayee

## Abstract

DNA double-strand breaks (DSBs) are a threat to genome stability. In all domains of life, DSBs are faithfully fixed via homologous recombination. Recombination requires the presence of an uncut copy of duplex DNA that is used as a template for repair. Alternatively, in the absence of a template, cells utilize error-prone Non-homologous end joining (NHEJ). Although ubiquitously found in eukaryotes, NHEJ is not universally present in bacteria. It is unclear as to why many prokaryotes lack this pathway. To understand what could have led to the current distribution of bacterial NHEJ, we carried out comparative genomics and phylogenetic analysis across ~6000 genomes. Our results show that this pathway is sporadically distributed across the phylogeny. Ancestral reconstruction further suggests that NHEJ was absent in the eubacterial ancestor, and can be acquired via specific routes. Integrating NHEJ occurrence data for archaea, we also find evidence for extensive horizontal exchange of NHEJ genes between the two kingdoms as well as across bacterial clades. The pattern of occurrence in bacteria is consistent with correlated evolution of NHEJ with key genome characteristics of genome size and growth rates; NHEJ presence is associated with large genome sizes and/or slow growth rates, with the former being the dominant correlate. Given the central role these traits play in determining the ability to carry out recombination, it is possible that the evolutionary history of bacterial NHEJ may have been shaped by requirement for efficient DSB repair.

## Introduction

Accurate transmission of genetic material from parent to progeny is essential for the continuity of life. However, low rates of error during replication and DNA break-inducing mutagenic agents (such as ionising radiation and reactive oxygen), while generating diversity for natural selection to act on (1), also adversely affect viability and could lead to diseases including cancer (2, 3). Therefore, most cellular life forms invest in mechanisms that repair damaged DNA including double-strand breaks (DSBs).

Two major mechanisms of repair of DNA double-strand breaks are homologous recombination and non-homologous end joining (NHEJ). Recombination-based repair requires a homologous copy of the DNA around the damage site for repair to occur. In contrast, NHEJ, the subject of the present work, directly ligates the double-strand break after detecting and binding the break ends (4, 5). Where direct ligation is not possible – at breaks that generate complex ends – a processing step involving the removal of damaged bases and resynthesis of lost DNA is required. Such processing can be error-prone (6). Thus NHEJ can be a double-edged sword: required for essential DNA repair when a homologous DNA copy is not available, but also prone to causing errors at complex DNA breaks.

NHEJ is a major mechanism of DNA repair in eukaryotes. In bacteria however, homologous recombination-based repair is most common; NHEJ, on the other hand, was described only recently (7, 8), and its prevalence still remains to be systematically elucidated. Unlike eukaryotes, where NHEJ activity is regulated in a cell-cycle dependent manner, it is unclear as to when NHEJ may be a preferred mode of bacterial DSB repair. Recent reports have shown that NHEJ can contribute to mutagenesis during a specific stage of bacterial growth, such as in stationary phase (5, 9), raising the possibility that availability of a second copy of the genome for repair and / or growth phase may dictate whether recombination or NHEJ is employed for repair.

Experiments in *Mycobacterium* and *Pseudomonas* have shown that bacterial NHEJ repair machinery consists primarily of two proteins. The homodimeric Ku binds DNA break ends and recruits the three- domain LigD harbouring phosphoesterase (PE), polymerase (POL) and ligase (LIG) activity. These three domains are respectively required to process the ends, add bases if necessary and subsequently ligate the break. Additionally, the POL domain mediates interaction between Ku and LigD (4, 10–12). As an exception, studies in *Bacillus subtilis* found that this bacteria encodes Ku along with a two- domain (LIG and POL) LigD (13). In the absence of LigD, it is also possible for LigC, which contains only the LIG domain, to carry out repair (4, 5, 14). Though *Escherichia coli* does not encode NHEJ, expression of M. tuberculosis Ku and LigD renders *E. coli* NHEJ proficient (15).

Studies in the early 2000s, with a small number of genomes, suggested that the distribution of NHEJ in bacteria could be patchy (16). Collectively, what does it mean for bacteria like *Escherichia coli* to not code for NHEJ and others like *Mycobacterium tuberculosis* to harbour it? Given the large population sizes and relatively short generation times that make selection particularly strong, the question of the pressures that determine the deployment of the potentially risky NHEJ assumes importance. In this study, using bioinformatic sequence searches of Ku and LigD domains in the genomic sequences of ~6000 bacteria, we have tried to 1) understand how pervasive their pattern of occurrence is, 2) trace their evolutionary history and, 3) understand what selection pressures could explain it.

## Methods

### Data

All genome information files for ~6000 bacteria were downloaded from the NCBI ftp website using inhouse scripts – whole genome sequences (.fna), protein coding nucleotide sequences (.fna), RNA sequences (.fna) and protein sequences (.faa). All the organisms were assigned respective phylum and subphylum based on the KEGG classification (https://www.genome.jp/kegg/genome.html).

### Identification of NHEJ repair proteins

Initial blastp (17) was run using each of the four protein domain sequences – Ku and LigD (LigD-LIG, LigD-POL and LigD-PE) from *Pseudomonas aeruginosa PAO1* as the query sequence against the UniProtKB database (18) with an E-value cut-off of 0.0001. The top 250 full length sequence hits were downloaded from Uniprot for both Ku and LigD domains respectively. A domain multiple sequence alignment was made using phmmer -A option with the top 250 hits as the sequence database and *P. aeruginosa* domain sequences as the query. An hmm profile was built using the hmmbuild command for the multiple sequence alignment obtained in the previous step. To find domain homologues, hmmsearch command with an E-value cut-off of 0.0001 was used with the hmm profile as the query against a database of 5973 bacterial genome sequences (Supplementary table 1). These homolog searches were done using HMMER package v3.3 (19). An in-house python script was written to extract the results and assign organisms with Ku and LigD domains. The same methods were used to identify NHEJ components in 243 archaeal genomes (Supplementary table 2).

Bacteria were assigned into five broad categories, based on the status of NHEJ components – *NHEJ-, Ku only, LigD only, conventional NHEJ+* and *non-conventional NHEJ+.* Conventional NHEJ was said to be present in bacteria harbouring Ku and LigD having LIG, POL and PE domains in the same protein. Non-conventional NHEJ included bacteria with Ku and at least one of the following: a) all LigD domains present in different combinations in different proteins, b) just the LIG and POL domains present in the same or, c) different proteins and, d) LIG with/without PE domain present in the same or different proteins. Organisms with PE and/or POL but not the LIG domain detected, were assigned to *NHEJ-* state if Ku domain wasn’t detected either; a *Ku only* state if Ku domain was detected (Figure 1).

**Figure 1:**
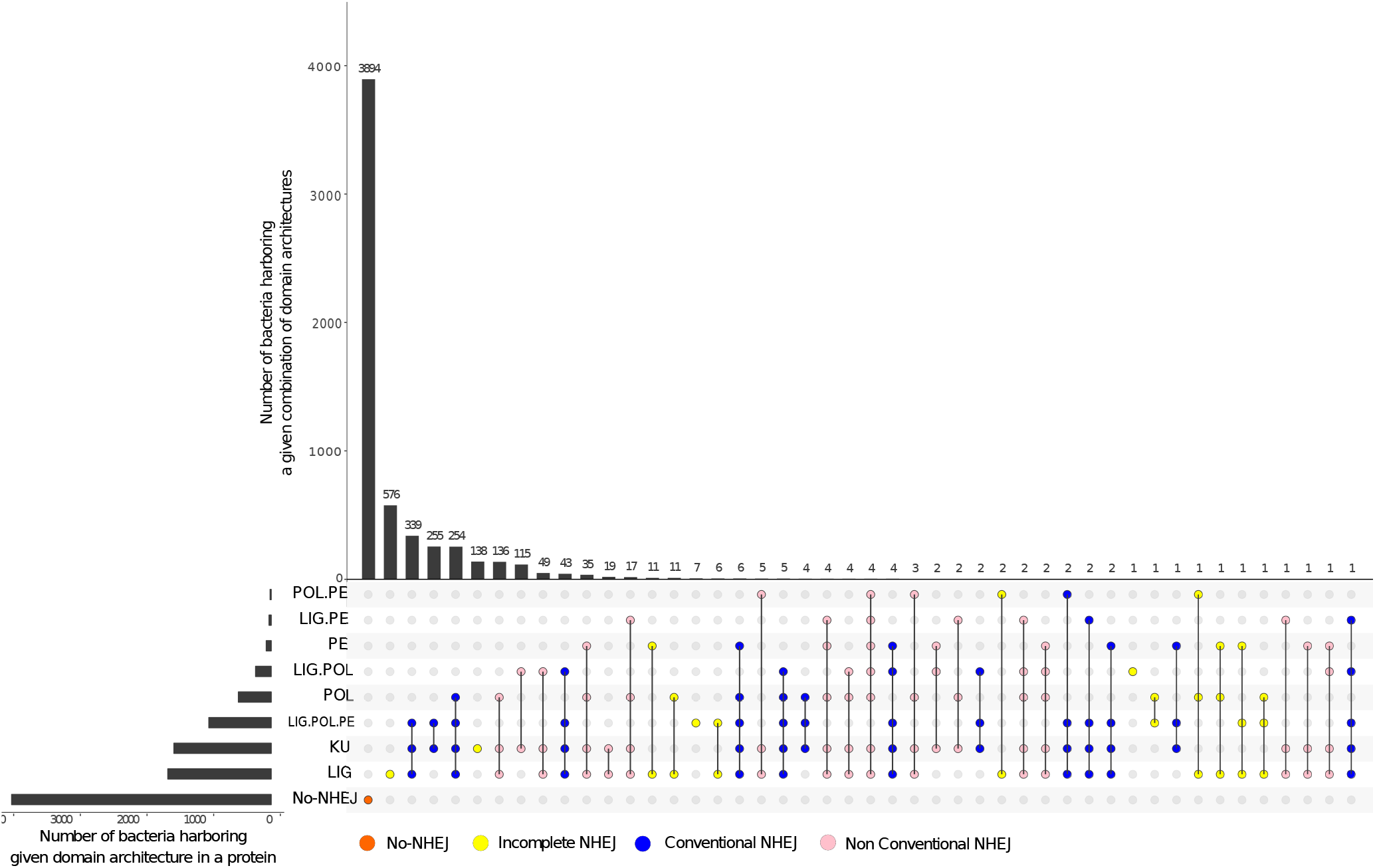
Distribution of NHEJ components in bacteria. An UpSet plot depicting the number of bacteria harboring a certain type of domain architecture per protein and number of bacteria harboring a combination of domain architecture in their respective genomes is shown for 5973 analysed genomes. NHEJ- (orange), Ku only (yellow), LigD only (yellow), Conventional NHEJ+ (blue) and non-conventional NHEJ+ (pink).

### Ku and LigD neighbourhood analysis

In-house python scripts were written to determine the proximity of *ku* and *ligD* on the genome using the annotation files. Taking one gene as the reference, the presence of the other gene was checked within a distance of 10 genes upstream or downstream inclusive of both strands. The organisation of NHEJ genes fell into three categories – 1) both genes on the same strand and within the distance range; this we call as operonic NHEJ, 2) both genes on different strands and within the distance range and 3) genes outside the distance range. This analysis was done only for *NHEJ+* organisms which code for LigD containing all the three domains as part of a single protein.

### Identification of rRNA sequences

A 16S rRNA sequence database was downloaded from the GRD (Genomic-based 16S ribosomal RNA database) website (https://metasystems.riken.jp/grd/). A multiple sequence alignment and a profile of the database was made using FAMSA v1.1 (20) and hmmbuild (19) respectively. To detect 16S rRNA homologues in our database of 5973 bacteria, nhmmer (21) was used with an E-value cut-off of 0.0001 where the GRD based 16S rRNA sequence profile was used as the query. In-house python script was written to parse the output files and select the best hits for further analysis.

### Calculation of genome sizes, growth rates and G-C content

In-house python scripts were written to calculate the genome sizes, growth rates and G-C content of all bacteria in our dataset (Supplementary table 1). These calculations were made after excluding the plasmid sequence information from the whole genome sequence assembly files. Previous studies have shown a significant positive correlation between a bacteria’s growth rate and the number of rRNA operons harboured in it’s genome (22–25). Therefore, rRNA copy number was taken as a proxy for growth rate.

### Genome size randomization analysis

For this analysis, 920 organisms harbouring conventional NHEJ were considered. The total number of coding DNA sequences (CDS) was used as the proxy for genome size. Since the chances of finding a gene in a larger genome are more than that in a smaller genome, 920 genes were drawn at random from a pool of all the genes coming from 5973 organisms in our dataset and every time the genome CDS size (genome size) from which the gene came from was noted. For each such iteration, the median genome size was calculated. This was repeated for 100 iterations and a distribution of median genome size was obtained. The non-parametric wilcoxon rank sum test was used to compare this random distribution to the median genome size of NHEJ harbouring bacteria.

### sHorizontal gene transfer analysis

Alien Hunter v1.7 (26) with default options was used to predict horizontally acquired regions (HARs) – based on their oligonucleotide composition – across bacterial genomes present in our dataset. In-house python scripts were used to detect Ku and LigD in the predicted HARs. NHEJ was said to be acquired through horizontal gene transfer if both Ku and LigD were present in the predicted HARs.

Phylogenetic incongruence between the archaeal-bacterial 16S rRNA based species tree and the corresponding gene trees (for Ku, LigD-LIG domain and RecA) was checked using the Approximately Unbiased (AU) statistical test (27). AU test was carried out with 10000 simulations with a constrained versus unconstrained approach (explained next). The multiple sequence alignments for each gene were run in an unconstrained mode for the given model parameters. Then the alignments were run in a constrained mode with respect to their respective species tree topologies. p-AU gives the probability of identifying the gene family as having evolved according to the species i.e the evolutionary history of the gene is the same as that of the species. This test was carried out using the -au options and -zb option implemented in IQTREE.

Further, to test the strength with which the aforementioned genes could be vertically transmitted or moved among closely or distantly related bacteria, non-parametric Mantel tests were carried out. Patristic distances were calculated between bacteria in the species tree and a given gene tree respectively. A Pearson product-moment correlation coefficient r was calculated between these two distance matrices consisting of bacteria shared between the two phylogenies under test. The significance of correlation was assessed via randomization test by conducting 10000 permutations of distance matrices. The correlation coefficient r was recalculated for each permutation to produce the null distribution and p-value was obtained using the one-tailed test using the R package.

RANGER-DTL 2.0 (28) was used to predict the donors and recipients of NHEJ horizontal gene transfer events among and between bacterial and archaeal species. A non-polytomous rooted 16S rRNA based tree with archaea as an outgroup was used as a species tree. Optimal rootings for Ku and LigD-LIG domain based bacterial-archaeal gene trees were determined using the *OptRoot* program with default options such that the Duplication-Transfer-Loss (DTL) reconciliation cost was minimized. *Ranger-DTL* program was used to compute the optimal DTL reconciliation of a given rooted species tree – rooted gene tree pair. In case of multiple optimal reconciliations. the program inherently reports an optimal reconciliation sampled uniformly at random. Therefore, the analysis was run for 100 simulations each with a transfer cost T=1, 2 or 3 (default) i.e a total of 300 simulations. The lower the transfer cost, the more the horizontal gene transfer events allowed during the reconciliations. *AggregateRanger* was used to compute support values for the most frequent mappings i.e the donor species, by accounting for the variance due to multiple optimal reconciliations and alternative event cost assignments. An in-house python script was written to back trace the most frequent recipient for a given most frequent mapping for both Ku (Supplementary table 3) and LIG domain (Supplementary table 4). The results were overlaid on the respective species phylogeny using the features provided in the iTOL server (https://itol.embl.de/login.cgi).

### Phylogenetic tree construction

For the construction of a species tree, one 16S rRNA sequence per genome was extracted into a multi-fasta file using an in-house python script. For bacteria with multiple 16S rRNA sequences, one 16S rRNA sequence was chosen such that it minimizes the number of Ns in that sequence and has the maximum sequence length. In order to build a pruned phylogenetic tree, 970 bacteria were randomly selected such that a genus was represented exactly once for every NHEJ state. Please note that in the case of non-conventional NHEJ, all its four sub-categories, as described in the previous sections, were treated separately, when including bacteria at the genus level. A multiple sequence alignment (MSA) was built using FAMSA 1.1 (20) with default options. The conserved regions relevant for phylogenetic inference were extracted from the MSA using BMGE v1.12 (29). After the manual detection of the alignment, one spurious sequence was removed and the MSA was built again for 969 bacteria (Supplementary table 5). Using IQTREE v1.6.5 (30), a maximum likelihood based phylogenetic tree was built with the best model chosen as SYM+R10 (LogL= −118152.5876, BIC= 249946.0604). ModelFinder (-m MF option) (31) was used to choose the best model for the tree construction compared against 285 other models. Branch supports were assessed using both 1000 ultrafast bootstrap approximations (-bb 1000 -bnni option) (32) and SH-like approximate likelihood ratio test (-alrt 1000 option) (33).

A similar approach was used to build a phylogeny of 1403 (Supplementary table 6) organisms, non-redundant at the species level, comprising just two NHEJ states- 1) *NHEJ-* and 2) *conventional NHEJ+.*

For the construction of bacterial-archaeal gene trees (Ku, LigD-LIG or RecA), only bacteria harbouring one Ku and one conventional LigD in their genomes were included. Since the motivation behind the construction of gene trees was to study the origin of NHEJ in bacteria, all archaea harbouring either Ku or LigD-LIG domain were included in the respective phylogeny. In species with multiple copies of RecA, RecA with the highest alignment length and identity was chosen for a given genome. The species in the bacterial-archaeal 16S rRNA based phylogeny depended on the species included in the corresponding gene tree. The approach used to build these phylogenies was similar to the one described in the previous two paragraphs.

### NHEJ ancestral state reconstruction analysis

To trace the evolutionary history, four discrete character states were defined as follows – *Ku only*, *LigD only, NHEJ-* and *NHEJ+.* States were estimated at each node using stochastic character mapping (34) with 1000 simulations provided as make.simmap() method in the R package phytools v0.6-44 (35). The phylogenetic tree was rooted using the midpoint method and polytomies were removed by assigning very small branch lengths (10^-6^) to all the branches with zero length. The prior distribution of the states were estimated at the root of the tree. Further, by default the method assumes that the transitions between different character states occur at equal rates. This might not always be true, especially with complex traits where it is supposedly easier to lose than gain such characters. Therefore, for the estimation of transition matrix Q, three discrete character evolution model fits were compared – Equal Rates (ER), Symmetric (SYM) and All Rates Different (ARD). This allowed for models that incorporate asymmetries in transition rates. Based on AIC weights, the ARD model (w-AIC_ARD_ =1, w- AIC_SYM_= 0, w-AIC_ER_= 0) was chosen as the best fit with unequal forward and backward rates for each character state transition. Finally, Q was sampled 1000 times from the posterior probability distribution of Q using MCMC and 1000 stochastic maps were simulated conditioned on each sampled value of Q. This strategy was used to reconstruct ancestral states for both phylogenies comprising 969 and 1403 organisms as described in the previous sections.

Ancestral states for genome size, a continuous trait, were reconstructed using fastAnc() method employed in phytools v0.6-44 based on the brownian motion model. This model was found to be the better fit model as compared to multiple rate model (BPIC_brownian_<BPIC_stable_ and PSRF approaching < 1.1 well within 1000000 iterations), assessed using StableTraits (36). The multiple rate model allows for the incorporation of neutrality and gradualism associated with Brownian motion and also includes occasional bursts of rapid evolutionary change. Ancestral states for growth rate were reconstructed by converting it into a binary trait. An organism was said to be slow growing if it encoded rRNA copy numbers less than or equal to the median rRNA copy number (median =3) and fast growing otherwise. The reconstruction was carried out using the same approach that was used for estimating NHEJ ancestral states, as described in the previous paragraph.

### NHEJ and genome characteristics phylogenetic comparative analysis

The two genome characteristics – genome size and growth rate – were compared across bacteria with different NHEJ states. The distributions across bacteria with different NHEJ states were first compared assuming statistical independence of bacteria, using Wilcoxon Rank sum test, wilcox.test() in R. Next, two measures of phylogenetic signal – Pagel’s λ (37) and Blomberg’s K (38) – were used for detecting the impact of shared ancestry, for genome size and growth rate across bacteria, using the phylosig() routine in phytools R package (35). A phylogenetic ANOVA, employed in phytools R package v0.6-44 (35), was carried out with 1000 simulations and Holm-Bonferonni correction to control for family-wise error rate, based on a method by Garland T. (39), to compare the genome characteristics in a phylogenetically controlled manner. Please note that genome sizes and rRNA copy numbers were log_i0_ transformed for all the analysis.

### Correlated evolution analysis

Two relationships – (NHEJ repair and genome size) and (NHEJ repair and growth rate) – were quantified using a statistical framework. To test if changes in genome characteristics occur independently of NHEJ or are these changes more (or less) likely to occur in lineages with (or without) NHEJ, two models of evolution were considered- independent and dependent. In the independent model, both the traits were allowed to evolve separately on a phylogenetic tree i.e non-correlated evolution. In the dependent model, the two traits were evolved in a non-agnostic manner i.e correlated evolution. The NHEJ repair trait had two repair character states – NHEJ- (0) and conventional NHEJ+ (1). Genome sizes, a continuous trait, were converted into binary state as well. For this, the mean genome size of organisms with 0 or 1 NHEJ repair state was computed. A “lower” state (0) was assigned if a value was less than the mean and a “higher” state (1) if the value was more. The same approach was used to convert growth rates (rRNA copy number) into a binary state.

A continuous-time Markov model approach was used to investigate correlated evolution between NHEJ repair and the genome characteristics. First, the maximum likelihood (ML) approach (40) was used to calculate log-likelihoods for the two models of evolution per trait pair – 1) NHEJ repair and genome size; 2) NHEJ repair and growth rate. A likelihood ratio statistic (LR) was calculated for both comparisons, followed by a Chi-square test to assess if the dependent model was a better fit. The degrees of freedom are given by: dfChi-square test = (n_rate-dependem model_ – n_rate-independent model_). There are 8 transition rates in the dependent model across four states (00,01,10,11) and four transition rates in the independent model across two states (0,1; 0,1). Therefore, the test was run with four degrees of freedom.

The ML approach implicitly assumes that the models used for hypothesis testing are free of errors. Therefore, to make the analysis robust, the bayesian rjmcmc (reverse jump markov chain monte carlo) approach was used to calculate the marginal log-likelihoods of the independent and dependent models of evolution (41). This approach takes into consideration the uncertainty and minimizes the error associated in calculating the parameters used in each of our model(s), ensuring reliable interpretations. log Bayes factor was used to assess the better fit out of the two models.

BayesTraits v3 (41) was used to carry out both ML and Bayesian rjmcmc based correlated evolution analysis as described above for both the models. ML was run using the default parameters. Bayesian rjmcmc was run for 5050000 iterations, sampling every 1000^th^ iteration with a burn-in of 50000. For the estimation of marginal likelihood, a stepping stone sampler algorithm was used where the number of stones were set to 100 and each stone was allowed to run for 10000 iterations.

### Phylogenetic logistic regression analysis

A method developed by Ives and Garland (42), was used to carry out the phylogenetic logistic regression analysis provided in the R package phylolm v2.6 (43) as the subroutine phyloglm() with method=logistic_IG10. NHEJ repair was the binary dependent variable with two states – NHEJ- (0) and conventional NHEJ+ (1). The two independent continuous variables were genome size (GS) and growth rate (GR). Three models were tested – 1) NHEJ ~ GS, 2) NHEJ ~ GR and 3) NHEJ ~ GS+GR. The best model was chosen according to AIC scores.

All the scripts used for analysis were written in python, perl or R. Statistical tests and data visualizations were carried out in R and iTOL server.

## Results

### NHEJ is sporadically distributed across bacteria

To identify NHEJ machinery across bacteria, we used the reference sequences of the Ku domain, and the LIG, POL and PE domains of LigD from *P. aeruginosa* to search ~6,000 complete bacterial genomes for homologs (see Methods). We defined bacteria encoding Ku and the complete, three- domain version of LigD as those harbouring a conventional NHEJ system. Organisms lacking the POL and/or the PE domains of LigD, and those encoding these domains in separate proteins, were defined as those carrying non-conventional NHEJ (Figure 1).

We found NHEJ in only ~1,300 (22%) genomes studied here. There were various combinations of Ku and LigD domains across these organisms, but a large majority (920) carried the conventional NHEJ. 75% bacteria harbouring conventional NHEJ coded for Ku and LigD in a 10kb vicinity of each other, with 60% organisms carrying Ku and LigD in an operon. Most bacteria (84%) harbouring NHEJ coded for a single copy of Ku, whereas the remaining coded for 2-8 Ku copies in their genomes. For example, as reported by McGovern et al. (44) and Kobayashi et. al. (45), we identified 4 Ku-encoding genes in *S. meliloti*. About two-thirds of NHEJ positive bacteria carried multiple copies of the LIG domain, 37% carried multiple copies of the POL domain and 8% bacteria had multiple copies of the PE domain (Supplementary table 1). We also noticed that 138 (2.3%) organisms encoded Ku and not LigD, and 619 (10.3%) only LigD and not Ku. (Figure 1; Supplementary table 1).

NHEJ was not restricted to specific bacterial classes (Figure 2), and was found in 10 classes. We found a significant enrichment of conventional NHEJ in Proteobacteria (Fisher’s Exact test, P = 3.8 x 10^-5^, odds ratio: 2.04) and Acidobacteria (Fisher’s Exact test, P = 5 x 10^-2^, odds ratio: 5.286). All Bacteroidetes with NHEJ harbour a conventional NHEJ, although we could not assign statistical significance to it (Fisher’s Exact test, P = 0.14, odds ratio: 1.38). In contrast, non-conventional NHEJ repair was significantly over-represented in Firmicutes (Fisher’s Exact test, P =1.23 x 10^-13^, odds ratio: 6) and Actinobacteria (Fisher’s Exact test, P = 2.12 x 10^-15^, odds ratio: 6.14). Twenty-four phyla did not include any NHEJ positive organism. (Supplementary table 7).

**Figure 2:**
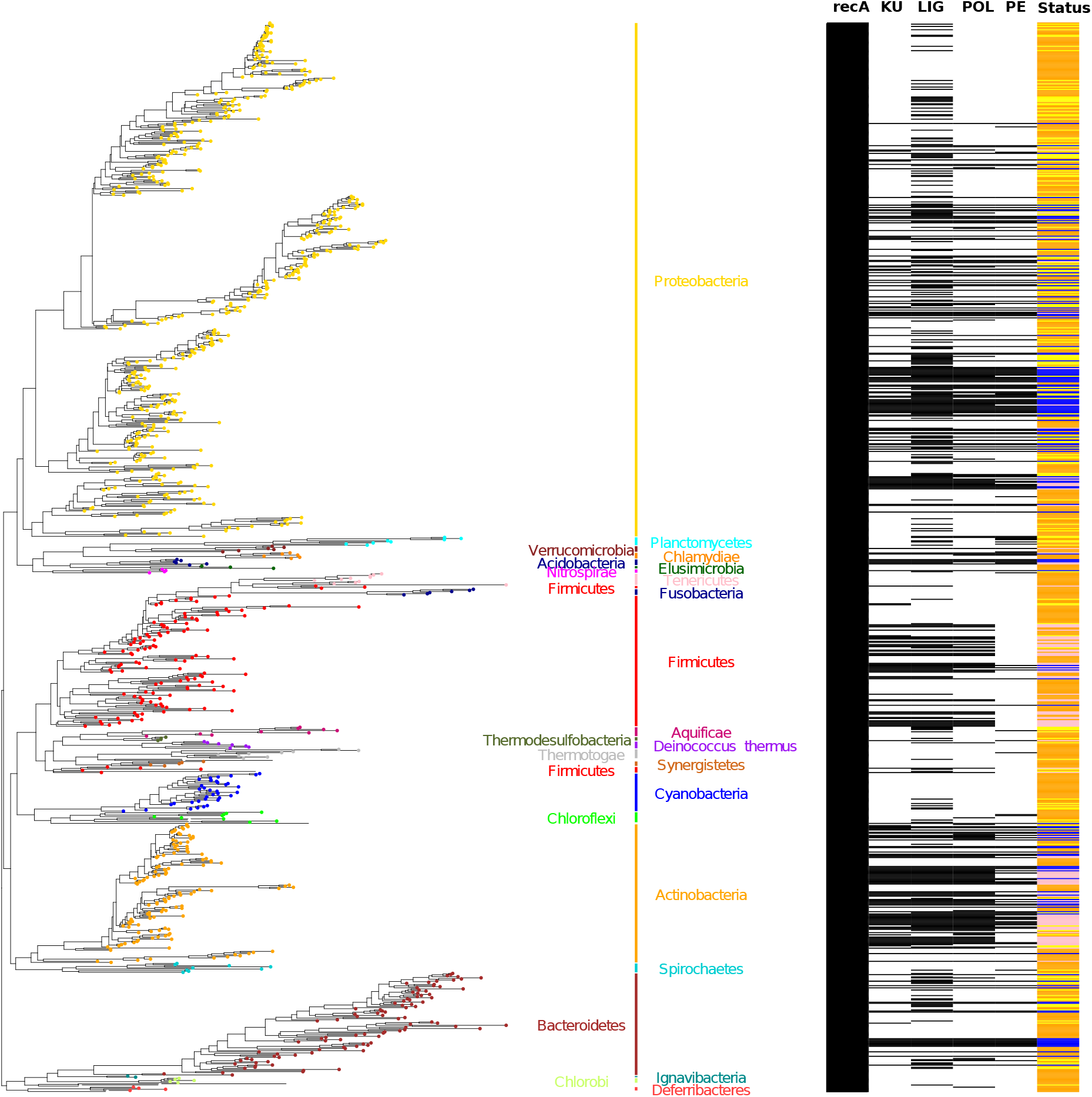
NHEJ is sporadically distributed across bacteria. 16S rRNA based species phylogenetic tree of 969 bacterial species (left) with presence/absence matrix of RecA, KU, LIG, POL and PE domains (right). Tip labels, representing bacteria, are coloured according to the phylum names and bars (right of the phylogenetic tree) that the given tip belongs to. The last column of the matrix represents NHEJ status corresponding to the species tip mapped to the phylogenetic tree. Status: Orange – NHEJ-; Yellow – Incomplete NHEJ; Blue – Conventional NHEJ+; Pink – Non-Conventional NHEJ+.

### NHEJ was gained and lost multiple times through evolution

We traced the number of NHEJ gains and losses starting from the eubacterial ancestor to the species at the tips of the 16S-based bacterial phylogenetic tree. To trace the evolutionary history we defined four discrete character states – *Ku only*, *LigD only* (conventional and non-conventional), *NHEJ-* and *NHEJ+.* Note that an *NHEJ+* state is defined only when both Ku and LigD are present in a bacterium. We calculated the posterior probabilities (pp) of each character state per node on the phylogeny, the distribution of the number of times each of the 12 character state transitions occurred (Figure 3A) and the distribution of the total time spent in each state (Figure S1). We performed this analysis for a set of 969 genomes in which each genus was represented once for each state (see Methods).

**Figure 3:**
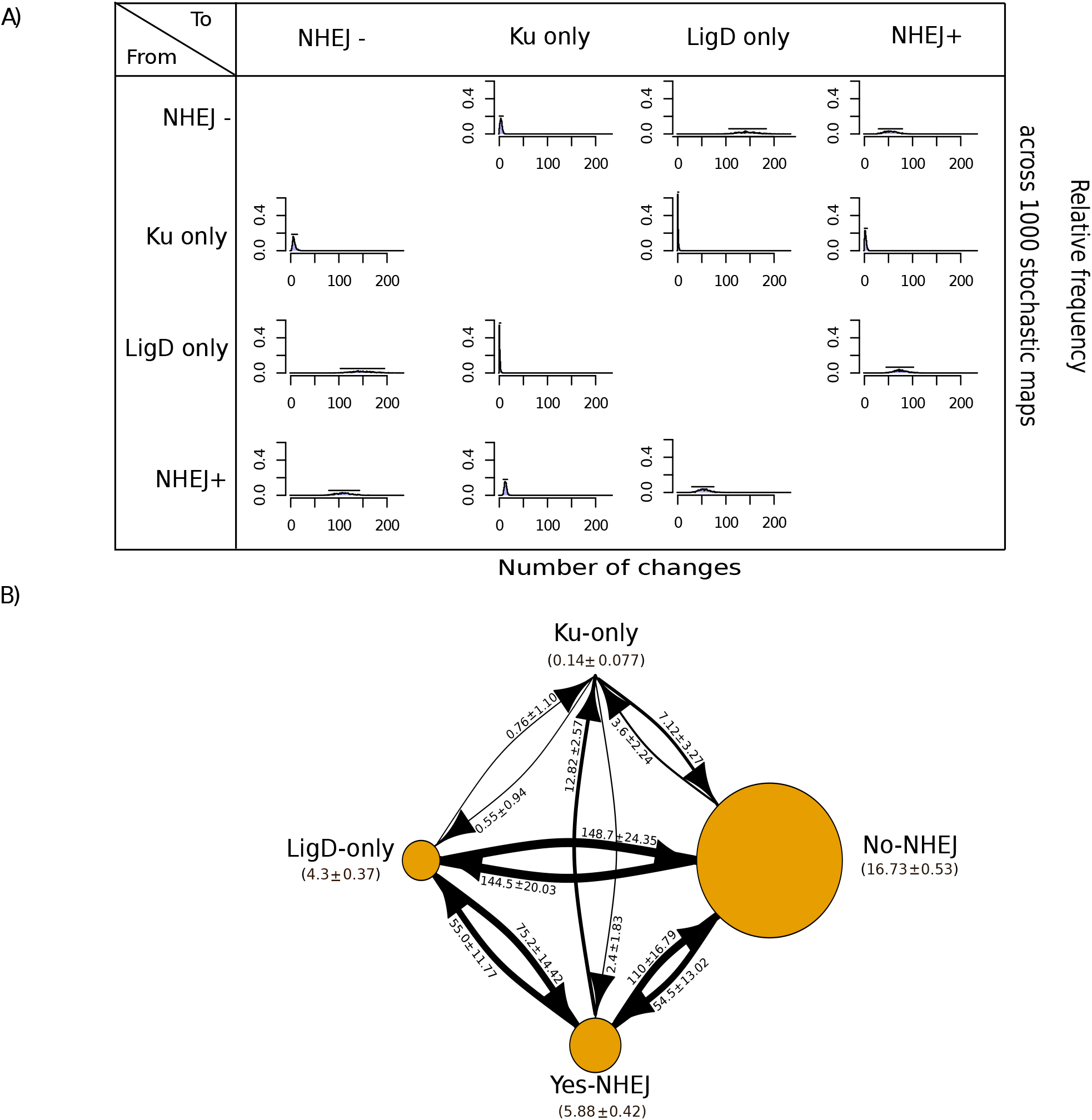
Transitions to a Ku-only state are rare. A) A matrix depicting relative frequency of number of changes of a state transition type across 1000 stochastic maps. B) A state transition diagram depicting the number of transitions between two given states and the time spent in each state during NHEJ evolution. The node size is proportional to the amount of time spent in a particular state. The arrow size is proportional to the number of transitions from one state to another.

We first asked if NHEJ was present in the common eubacterial ancestor and, given the sporadicity of NHEJ, subsequently lost in several lineages (Supplementary table 8). We assigned a major primary gain to an internal ancestral node if (a) all nodes leading to it from the root had *NHEJ-* state; (b) the pp of either *NHEJ+, LigD only* or *Ku only* at that ancestral node was >=0.7; (c) a gain of *LigD only* or *Ku only* was followed by a transition to *NHEJ+* and; (d) if it had at least three descendent species. We observed multiple major independent primary gains at ancestral nodes within Bacteroidetes, Actinobacteria, Firmicutes, Acidobacteria and multiple sub-clades of Proteobacteria (Supplementary table 8). It follows that the common eubacterial ancestor likely did not have NHEJ (Figure 4).

**Figure 4:**
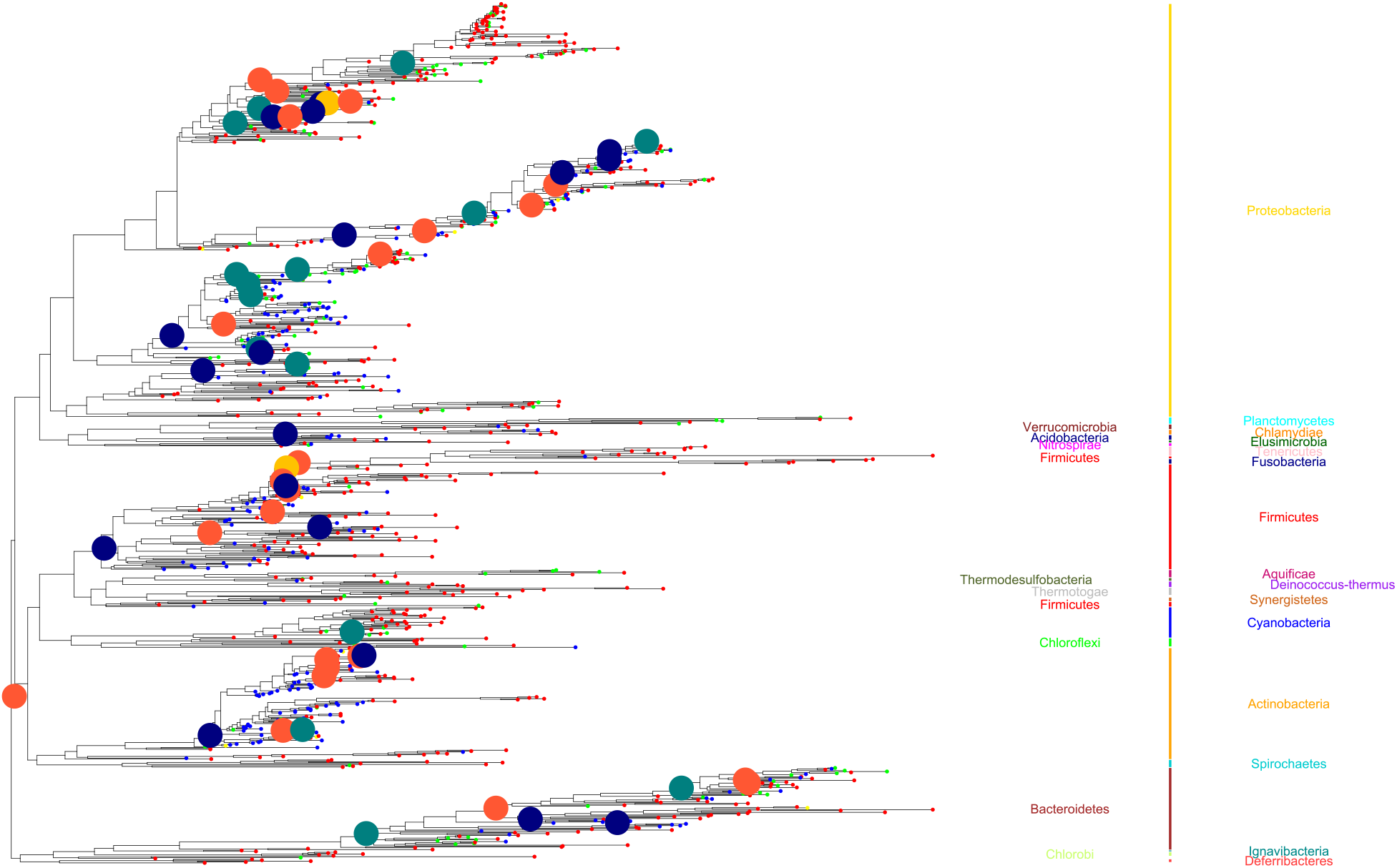
NHEJ was gained and lost multiple times through evolution. A trace of the evolutionary history of the two-component NHEJ system across bacteria. The tip and node labels are coloured according to the NHEJ states: Red – NHEJ-; Yellow – Ku only; Green – LigD only; Blue – NHEJ+ (Conventional and Non-conventional). NHEJ state for nodes is shown only when the posterior probability support is greater than 70%; interpreted as change in NHEJ state at that node as compared to shallower phylogenetic depths.

The gain of NHEJ can be sequential, gaining either *Ku only* or *LigD only* followed by the gain of the other component; or it can be a one-step acquisition of both components (Figure 3B). The most common transition from an NHEJ- state was to a LigD only state. Also frequent was the direct acquisition of both components to transition from an *NHEJ-* to an *NHEJ+* state. Transition from *NHEJ-* to *Ku only* was negligible. In the reverse direction, a one-step loss of both Ku and LigD is the most likely. Again, *Ku only* state is rare.

A one-step transition from *NHEJ-* to *NHEJ+* is likely through horizontal gene transfer. 60 bacterial genomes belonging to the phyla Alpha-proteobacteria (in particular the Rhizobiales) – and Beta- proteobacteria, and Streptomycetales carried their NHEJ components on plasmids (Supplementary table 1). However, based on abnormal word usage statistics (see Methods), we couldn’t find NHEJ to be a part of the horizontally acquired component of the chromosomes of any bacterial genome. At least two *NHEJ-* to *NHEJ+* transitions occurred close to the root, and it is possible that the predictions of horizontally acquired NHEJ systems made so far may be an underestimate (Figure 4). We investigate this in greater detail below.

In summary, (a) the common eubacterial ancestor was devoid of NHEJ; (b) NHEJ was gained and lost multiple times; (c) transitions to a *Ku only* state are rare.

### NHEJ and horizontal gene transfer

Experimental studies in the archaea *Methanocella paludicola* (46–48) have confirmed the presence of a functional NHEJ repair, with crystal structures revealing close relationship with the bacterial proteins (47, 49).

To assess the possibility of horizontal transfer of NHEJ machinery across prokaryotes, we first performed a domain wise search in 243 archaea. These searches revealed the presence of both Ku and LigD-LIG domains in 10 archaeal species (see Methods; Supplementary table 2). However 230 archaeal genomes encoded LigD but no Ku. This lends support to previous reports suggesting that NHEJ is rare in archaea (46, 47, 49).

In order to check for horizontal transfer events between bacteria and archaea, we used phylogenetic methods based on detecting conflicts between an organismal phylogeny and a phylogeny inferred for Ku and LigD-LIG domains respectively. This method allowed us to test for any ancient transfers across bacteria as well. We found that NHEJ proteins undergo horizontal gene transfer events at significantly high rate (p-AU=0; Figure 5B AND 5C; see Methods) and these are not limited to closely-related species or co-speciation events (Figure 5D; Mantel test, P < 10^-4^, r_Ku_=0.25; P < 10^-4^, r_LIG_= 0.4). We also noted incongruence with respect to the RecA phylogeny (p-AU=0; Figure 5A) as has been reported before (50, 51). However, these were limited at best to transfers among closely related species (Figure 5D; Mantel test, P < 10^-4^, r_RecA_ 0. 94).

**Figure 5:**
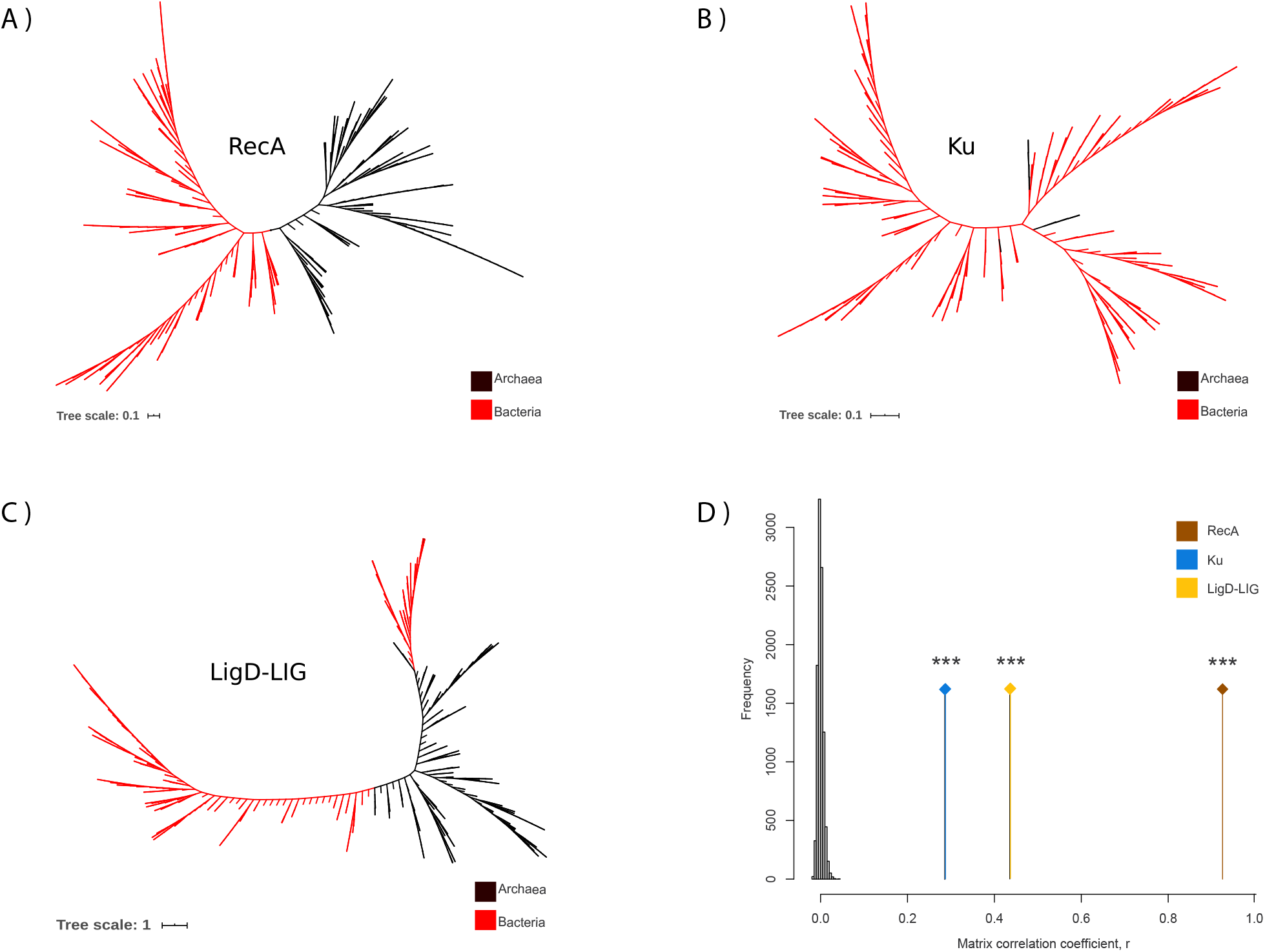
Phylogenetic methods suggest a strong role of Horizontal Gene Transfer in NHEJ evolution. A) Unrooted RecA tree, B) Unrooted Ku tree, C) Unrooted LigD-LIG tree and, D) Mantel test correlation coefficient (r) comparing RecA, Ku and LigD-LIG distance matrices with 16S rRNA distance matrices respectively, compared against a null distribution of r obtained by 10000 matrix randomizations.

An incongruence between a species and gene tree could result due to processes other than horizontal gene transfer, like duplications and losses. Therefore, to predict the most frequent transfer events, we used a reconciliation approach based on the duplication-transfer-loss (DTL) model. DTL employs a parsimonious framework where each evolutionary event is assigned a cost and the goal is to find a reconciliation (possible evolutionary history of gene tree inside a species tree) with minimum total cost. We observed a high rate of horizontal gene transfer between bacterial clades – Firmicutes, Actinobacteria, and Proteobacteria and Archaea, where each of these played the role of a donor and a recipient in Ku (Figure 6A) and LigD-LIG transfer events (Figure 6B). We observed that all proteobacterial species- with the exception of delta-proteobacteria, which was a donor of Ku to archaeal recipients – were recipients of both Ku and LigD from archaea or other distantly related bacteria. On one hand, we found strong evidence of Ku transfers from Archaea to Firmicutes and Actinobacteria and on the other hand, LigD-LIG transfers most likely occurred from Firmicutes and Actinobacteria to Archaea. Together, this raises the possibility of NHEJ transfers between bacteria and archaea.

**Figure 6:**
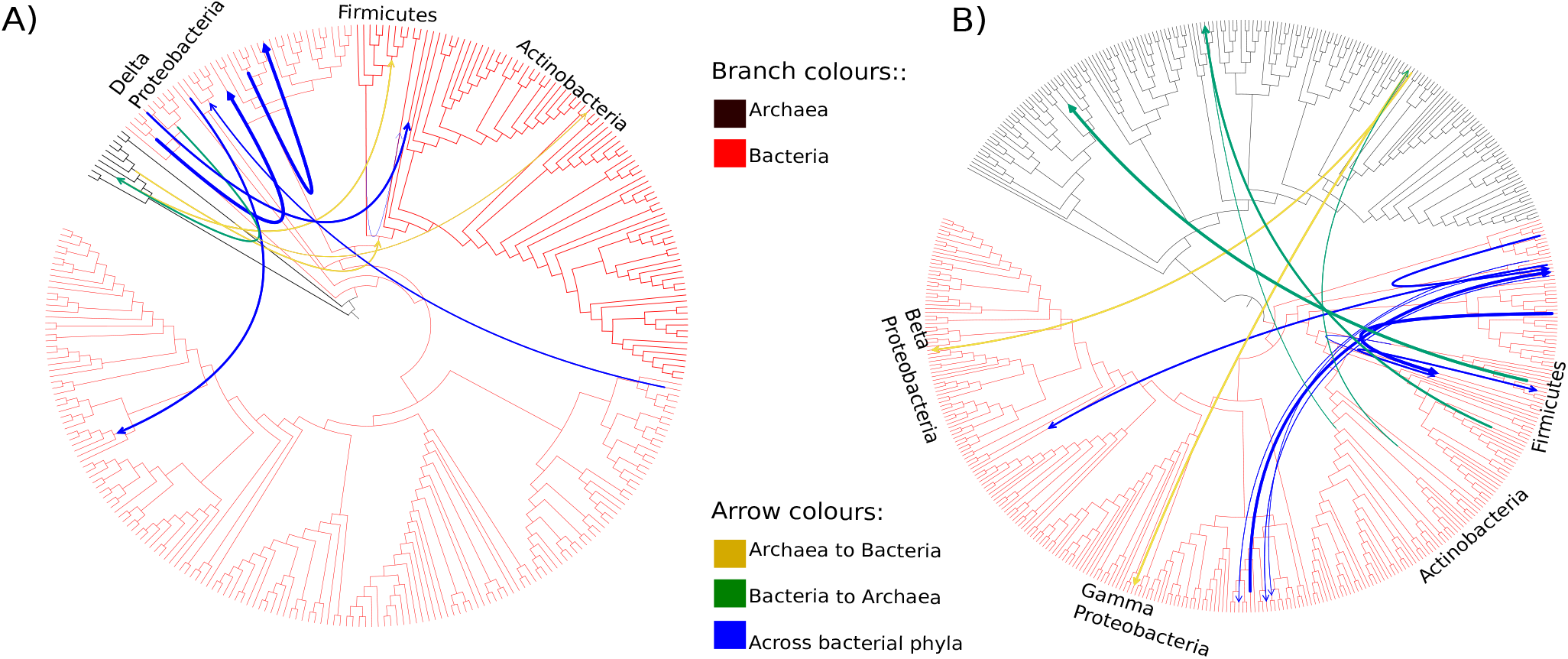
Extensive HGT among bacterial phyla and between bacteria and archaea. Species tree depicting the most frequent donor-recipient pairs involved in A) Ku HGT events, B) LigD-LIG HGT events. The width of the arrow corresponds to the number of reconciliations supporting a given transfer event.

### NHEJ occurrence is associated with genome size, growth rate and G-C content

Recently, Ku-encoding organisms were shown to have higher genomic G+C content (52). Given its central role in DNA repair, we asked whether any other genome characteristics could also be associated with the presence or absence of NHEJ. First, we verified that the findings of Weissman et al. on the correlation between the presence of Ku and G+C content held true for NHEJ+ states as defined in our study (Figure 7A, Supplementary S4-S5, S7C) (52). Along with this, we tested two additional characteristics: genome size and growth rate (as measured by the copy number of rRNA operons), both of which could determine the availability or the lack of a homologous template for high fidelity recombination-based repair. We restricted these analyses to conventional NHEJ-harbouring bacteria as a proxy for repair proficiency, and compared them to *NHEJ-* genomes. Data including non- conventional NHEJ are shown in Supplementary Figures S2 and S3.

**Figure 7:**
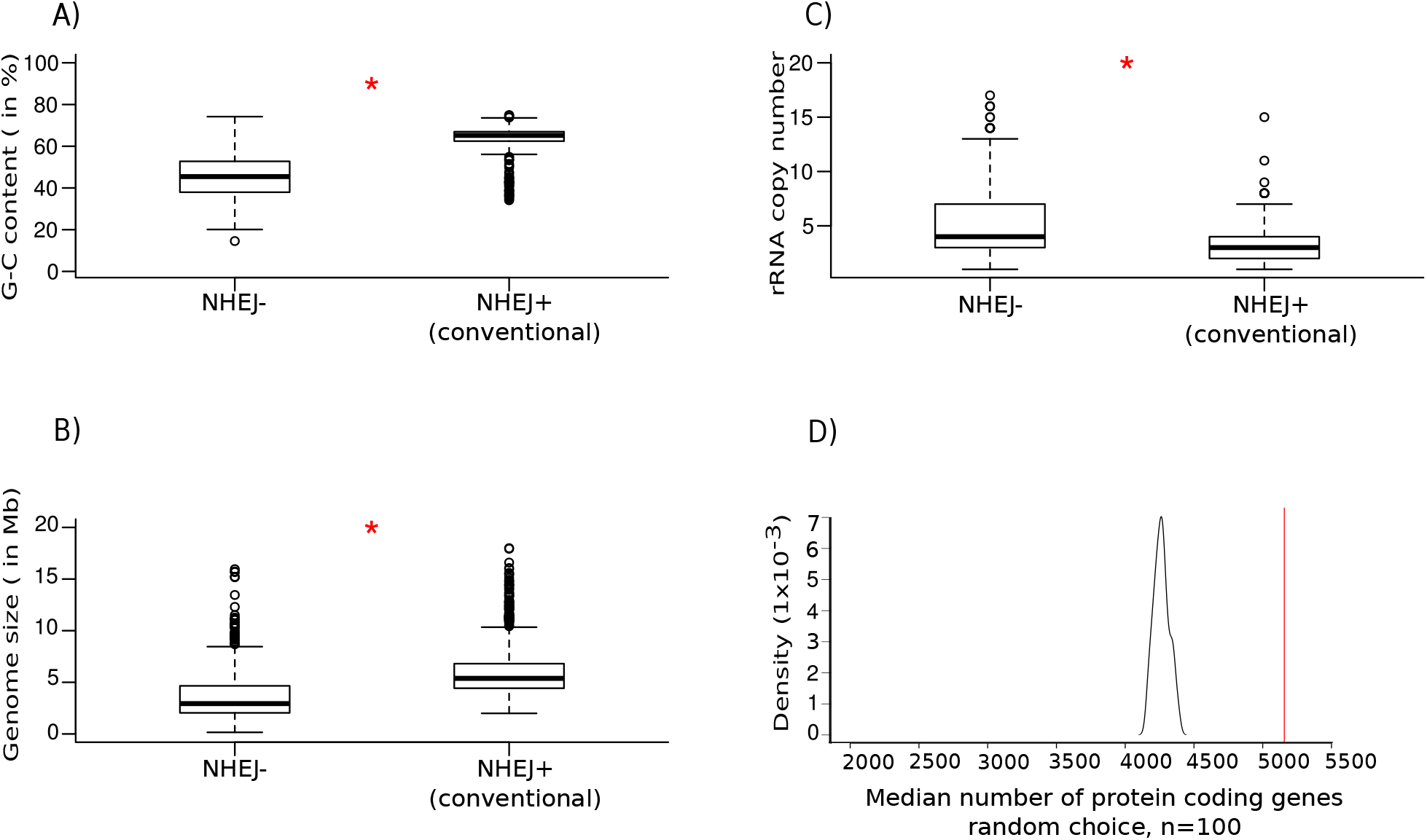
NHEJ presence and absence is associated with genome size, growth rate and G-C content. A) Boxplot comparing the distribution of G-C content between NHEJ– and conventional NHEJ+ bacteria. B) Boxplot comparing the distribution of genome size between NHEJ- and conventional NHEJ+ bacteria. C) Boxplot comparing the distribution of rRNA copy number between NHEJ- and conventional NHEJ+ bacteria. D) A density distribution plot depicting the distribution of genome size (median number of protein coding sequences) expected by a random distribution (black) where the probability of having NHEJ is linearly proportional to genome size and the median genome size of organisms harboring NHEJ (red).

Bacteria with NHEJ were found to have larger genomes (median = 5.4 Mb) than those without NHEJ (Median= 2.9 Mb; Wilcoxon rank-sum test, P < 10^-15^; Figure 7B, S7A), and significantly larger than that expected by a random distribution in which the probability of having NHEJ is linearly proportional to genome size (Wilcoxon rank sum test, P < 10^-15^, across 100 simulations; Figure 7D). This relationship was found to be true within the phylum Proteobacteria, Actinobacteria, Bacteroidetes and Firmicutes as well (Supplementary figure S6).

In addition, bacteria harbouring NHEJ were found to have significantly fewer rRNA copies (median = 3), and by inference slower growth rates, than bacteria without NHEJ (median = 4; Wilcoxon rank-sum test; P < 10^-15^; Figure 7C). While the distribution of rRNA copy numbers for genomes without NHEJ was broad, those with conventional NHEJ fell within a narrow range, representing relatively slower growth (Supplementary figure S7B). At the phyla level, this relationship was found to hold true for Proteobacteria and Actinobacteria, whereas there was no significant difference for Bacteroidetes and Firmicutes respectively (Supplementary figure S8).

In order to confirm the result in a phylogenetically controlled manner, Pagel’s λ and Blomberg’s K were used to first measure whether closely related bacteria tended to have similar genome sizes and growth rates in the dataset (see Methods). These measures suggest that phylogenetic coherence is significantly greater than random expectations for both the genome characteristics (Supplementary Table 9). Therefore, the distributions of genome size between bacteria with different NHEJ status were compared while accounting for the statistical non-independence of closely related taxa (see Methods). We found a significant difference in log_10_(genome sizes) between bacteria with conventional NHEJ and without the repair (phyloANOVA; P = 6 x 10^-3^); the characters being mapped on the phylogenetic tree of 969 bacteria with five discrete groups – *NHEJ-, Ku only, LigD-only, conventional NHEJ+ and non-* Table 1 *conventional NHEJ+.* However, we did not observe a significant difference in log_10_ (rRNA copy number) between the two groups of bacteria (phyloANOVA; P = 1; See Discussion).

**Table 1:**
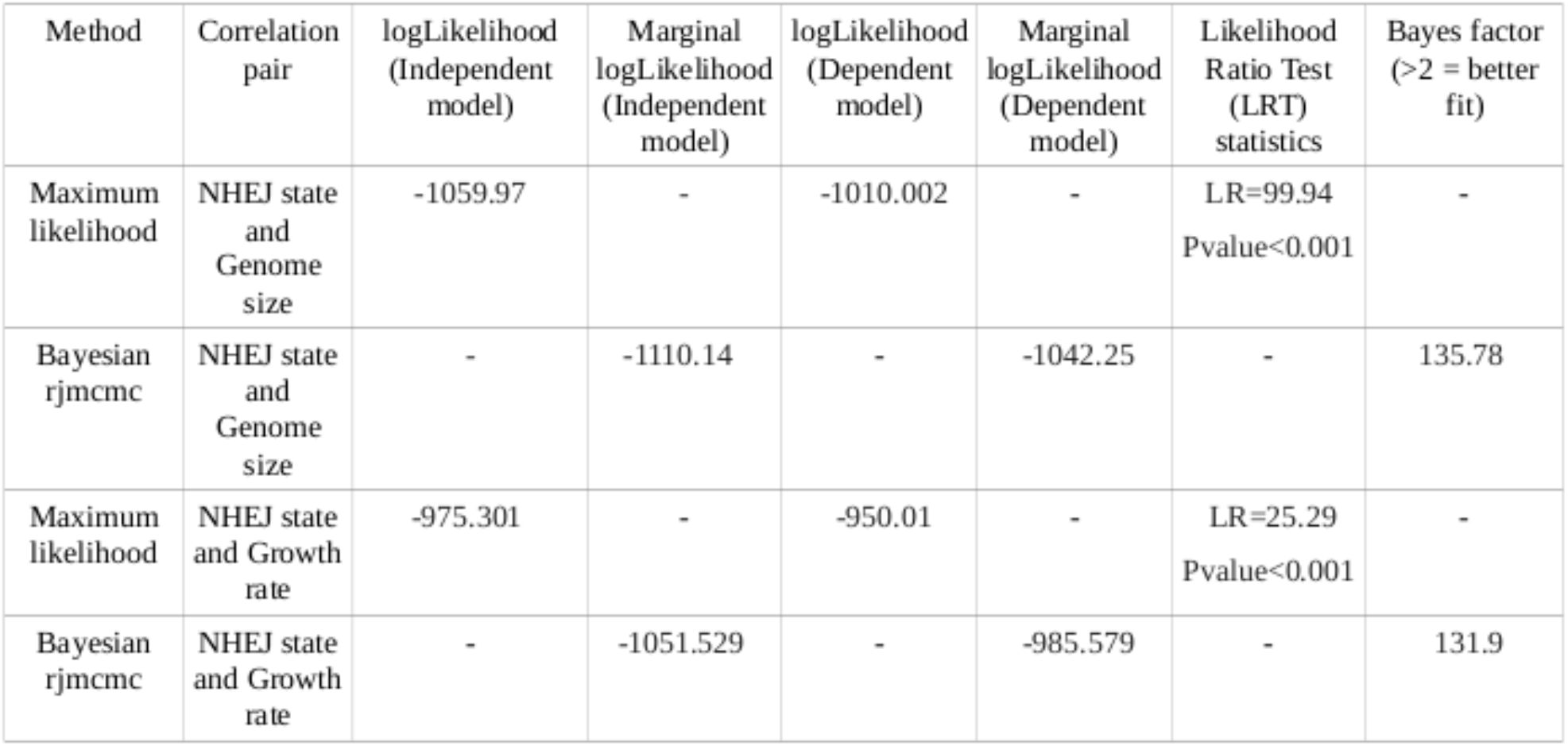

We used maximum likelihood as well as Bayesian approaches to test if the observed associations of conventional NHEJ individually with large genomes and slow growth rates is indicative of dependent or independent evolution of these traits on the phylogenetic tree (see Methods). Both suggested that the phylogenetic data fit models of evolution in which conventional NHEJ presence or absence and genome size or growth rate are evolving in a correlated manner (Table 1). This strengthens the association between the tested variables in a phylogenetically controlled way. Phylogenetic logistic regression of the conventional NHEJ occurrence with both the continuous independent variables showed, however, that genome size is the stronger correlate (see Methods; Table 2).

**Table 2:**
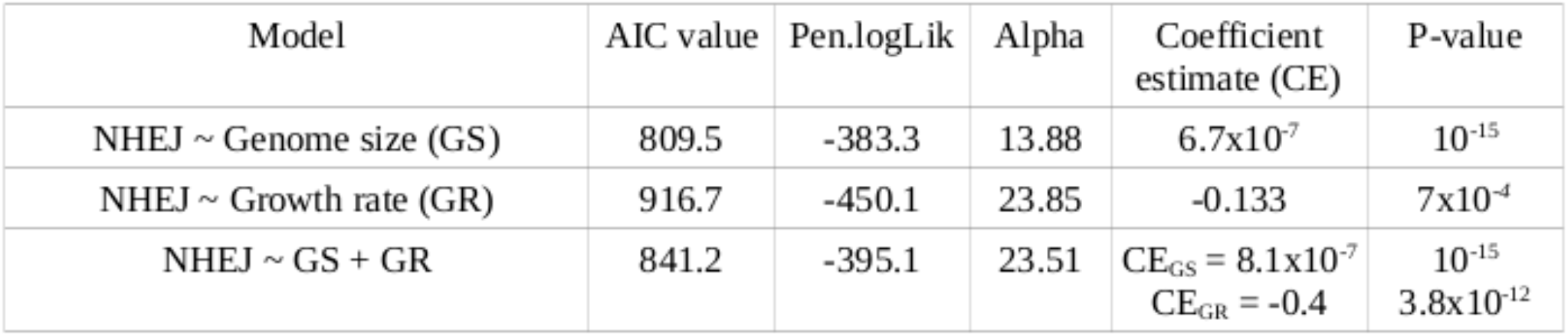

| Model | AIC value | Pen.logLik | Alpha | Coefficient estimate (CE) | P-value |
| --- | --- | --- | --- | --- | --- |
| NHEJ ~ Genome size (GS) | 809.5 | -383.3 | 13.88 | 6.7x10<sup>-7</sup> | 10<sup>-15</sup> |
| NHEJ ~ Growth rate (GR) | 916.7 | -450.1 | 23.85 | -0.133 | 7x10<sup>-4</sup> |
| NHEJ ~ GS + GR | 841.2 | -395.1 | 23.51 | CE<sub>GS</sub> = 8.1x10<sup>-7</sup><br>CE<sub>GR</sub> = -0.4 | 10<sup>-15</sup><br>3.8x10<sup>-12</sup> |

As a case study where the association of both genome size and growth rate with NHEJ evolution was prominent, we found a gain of conventional NHEJ in the ancestor of two genera belonging to Corynebacteriales – *Mycobacterium* and *Corynebacterium* – where the former retained and the latter had a secondary loss of the machinery. Phylogenetic ancestral reconstruction analysis revealed an increase in genome size in the ancestor of Corynebacteriales, followed by an NHEJ gain. While *Mycobacterium* retained NHEJ, *Corynebacterium* lost the machinery along with a decrease in genome size. Using a similar analysis, growth rate mapped to this sub-clade revealed an increase in rRNA copy number in *Corynebacterium*. (Figure S9; see methods).

## Discussion

Taking advantage of the availability of a large number of genomes, we confirm the non-ubiquitous nature of NHEJ across the bacterial domain (7, 14) with 22% bacteria coding for it. At the taxa level, while some phyla retain this sporadicity, others are devoid of NHEJ repair machinery, consistent with previous studies on distribution of bacterial NHEJ proteins (14). In line with previous reports on the multiplicity of NHEJ in certain bacteria (6, 44, 53–56), we further discovered a sporadic presence of multiple copies of NHEJ proteins extending to different subclades of Proteobacteria, Actinobacteria, Firmicutes, Bacteroidetes etc. (Supplementary table 1). A trace of the evolutionary history suggested that NHEJ was most likely absent in the eubacterial ancestor. Furthermore, 96% archaeal genomes coded for LIG but no Ku. Therefore, by inference, NHEJ was absent in the Most Recent Common Ancestor (MRCA) of all prokaryotes as well. It was instead gained independently multiple times in different bacterial lineages. However, these primary gains were not sufficient to stabilize the repair states in sub-clades, since a large number of secondary losses and gains were observed across the phylogeny (elaborated further below).

Our analysis suggests that there are two common methods to arrive at an *NHEJ+* state- a) the most common way to gain NHEJ was by the acquisition of LigD followed by Ku and b) a direct transition from an *NHEJ-* to *NHEJ+* state. We found that the LigD only state is more prevalent and is a prominent intermediate in the evolution of the *NHEJ+* state in Proteobacteria and Bacteroidetes (pp_Proteobacterial-subclade_ 0.875 and pp_Bacteroidetes_=0.7; Figure S10C and S10D). However, 90% of *LigD only* genomes did not encode POL and PE domains, raising the possibility that *LigD only* to *NHEJ+* transitions could actually be *NHEJ-* to *NHEJ+* transitions.

Direct NHEJ gain can be explained by their acquisition via horizontal gene transfer (HGT). In this direction, we observed 60 *NHEJ+* organisms coding this repair on their plasmids. However, our analysis based on the detection of base compositional differences revealed no horizontally acquired chromosomal NHEJ. These numbers may be an underestimate because one is unlikely to pick HGT events if – (a) the regions getting horizontally transferred have the same compositional biases in the donor and the recipient cells and/or (b) such HGT events occur early in bacterial evolution. Two instances of the latter case that we observe in our analysis are direct primary gains of NHEJ at the ancestral nodes of Firmicutes and Actinobacteria, with high posterior probability support (pp_Firmicutes_=0.987 and pp_Actinobacteria_ = 0.967; Figure S10A and S10B). Aravind and Koonin (7), proposed a possibility of HGT between bacteria and archaea. In this direction, using a gene tree-species tree reconciliation approach, we observed evidence for HGT between bacteria and archaea. Furthermore, this method allowed us to find evidence of HGT between distantly related bacteria as well (Supplementary table 3 and 4). It should be noted that for the lack of methods to accurately date bacterial species trees and because transfers from unsampled species or extinct lineages could violate time constraints, we used an undated phylogeny for this analysis. Therefore, for each most frequent donor predicted, we noted the most frequent recipient as the potential donor-recipient pair. A similar search in ~3000 actinobacteriophages yielded no hits, although, Pitcher et. al. (57) have experimentally shown the employment of *M. smegmatis* LigD by the Ku expressing mycobacteriophages Omega and Corndog for a successful infection. Thus, while our results suggest no transfer events between bacteria and bacteriophages, whether this is an artefact of sequence amelioration processes (58), remains to be tested.

A third route to an *NHEJ+* state could have been via a *Ku only* state. However, we observed that the time spent in a *Ku only* state is the least and NHEJ gains via this route are rare. Approximately 90% *Ku only* states are present in Firmicutes alone, specifically belonging to two genera, *Bacillus* and *Fictibacillus*. This suggests that *Ku only* state is largely avoided across most bacteria. This could be because Ku alone is non-functional in repair or in some cases could even block the access of break ends for recombination-based repair (59–61). In 138 organisms, where Ku is retained in the absence of LigD, it’s function could be mediated by cross-talk with other ligases such as LigA. Consistent with this, it has been previously shown that if damage produces 3’ overhangs specifically, LigA could repair the lesion even in the absence of LigB/C/D (4). *In vitro* studies have also shown that *M. smegmatis* Ku can stimulate T4 DNA ligase (62). We observed the same trend in Archaea; Ku was always present with a LIG domain in 10 archaeal species, whereas 230 archaea coded for LIG domain and no Ku. These observations are consistent with experimental studies reporting a fully functional complement of NHEJ being present only in a single archaea, and with the likelihood of microhomology-mediated end joining (via ligase alone) being more prevalent in these organisms (46, 49).

It is likely that several factors have contributed to the current distribution of NHEJ in bacterial systems. For example: 1) Studies in both prokaryotes and eukaryotes have suggested a cross-talk between NHEJ and other repair mechanisms like recombination-based repair (55, 56, 63, 64), base excision repair (64) and mismatch repair (65). Shen et. al. (65) have shown that a bacteriophage infection is prevented by generating DSBs produced by MutL, a conventional mismatch repair endonuclease, which is subsequently repaired by NHEJ via a large deletion. However, this might not be a conserved pathway of repair as our analysis suggests an absence of NHEJ in majority of actinomycetes belonging to the genus Corynebacterium including *C. glutamicum* and 96% of archaea. 2) NHEJ could co-occur with other pathways that may or may not be directly involved in DSB repair. For example, NHEJ has been found to be active during sporulation in *Bacillus subtilis* (64, 66), however, non-spore-forming bacteria, also encode for NHEJ (7, 44 and our work). Additional cross-talk at the level of regulation has also been reported, such as in *Mycobacterium*, where the deacetylase Sir2 has been implicated in regulating NHEJ (67). It is equally plausible that signatures of displacement of certain domains from genomes encoding NHEJ could inform us on how acquisition of NHEJ could have shaped genome architecture. In line with this, we did note 20 additional domains coded in Ku and LigD proteins (Supplementary table 10) across different phyla. These would be useful to experimentally test in the future to understand whether they show functional interaction with NHEJ repair.

To understand the selection pressures that might play a role in shaping the evolutionary pattern of NHEJ, we focussed our efforts on studying the genome characteristics associated with this DNA repair. We reasoned that under the event of a DSB, the unavailability of a template would prevent the highly accurate homologous recombination repair to act. Thus, factors that affect rates of genome duplication or probability of multiple DSBs occurring could be central to determining the need for NHEJ-based repair. Therefore, one might expect a higher selection pressure of maintaining NHEJ in bacteria with slower growth rates and larger genome sizes. In line with our hypothesis, we found that the organisms possessing conventional NHEJ tend to have significantly larger genome size and slower growth rate as compared to those that are devoid of it. A phylogenetically controlled test suggested that this association held true for genome size and not growth rate. We note that phyloANOVA used here for hypothesis testing assumes normality. However, our data for genome size and rRNA copy number are not normally distributed, even after converting them into a logarithmic scale (Shapiro-Wilk normality test; P_genome size_ = 1.7 x 10-10, P_rRNA copy number_ < 10-15). Therefore, the result should be interpreted keeping this caveat in mind. Furthermore, Maximum Likelihood and Bayesian inferences showed that there is a correlated evolution of NHEJ with genome size and growth rate along the phylogenetic tree. Considering the two variables together, we found that genome size is the better correlate of the *NHEJ+* state. Therefore, we conclude that the *NHEJ+* state is strongly associated with genome size and to a much smaller extent with growth rate throughout its evolution.

In sum, our study highlights the evolutionary trajectory of NHEJ and central characteristics that may have determined it’s sporadic distribution. DSB repair, including NHEJ, has been implicated in shaping bacterial genomes through mutagenesis (9), HGT (68) and their effect on genomic GC content (52). Given this relationship between repair and genome evolution, it is important to ask how one factor may have influenced the other during bacterial evolution. It is also possible that similar forces have played a role in the evolution of other repair pathways and the genomes encoding them.

## Abbreviations

NHEJ: Non Homologous End Joining,
DSB: Double Strand Break,
DNA: Deoxyribonucleic Acid,
rRNA: ribosomal Ribonucleic Acid,
LIG: Ligase domain,
POL: Polymerase domain,
PE: Phosphoesterase domain,
CDS: Coding DNA sequence,
HAR: Horizontally Acquired Region,
HGT: Horizontal Gene Transfer,
AIC: Akaike Information Criterion,
BPIC: Bayesian Predictive Information Criterion,
PSRF: Proportional Scale Reduction Factor,
ML: Maximum Likelihood,
RJMCMC: Reverse Jump Markov Chain Monte Carlo,
GS: Genome size,
GR: Growth Rate

## Acknowledgment

We thank Saurabh Mahajan for discussions. We thank Supriya Khedkar and members of the AB and AS labs for comments on the manuscript.

