## Supplementary_figures for "Evolutionary and Comparative analysis of bacterial Non-Homologous End Joining Repair"

**Figure S1**

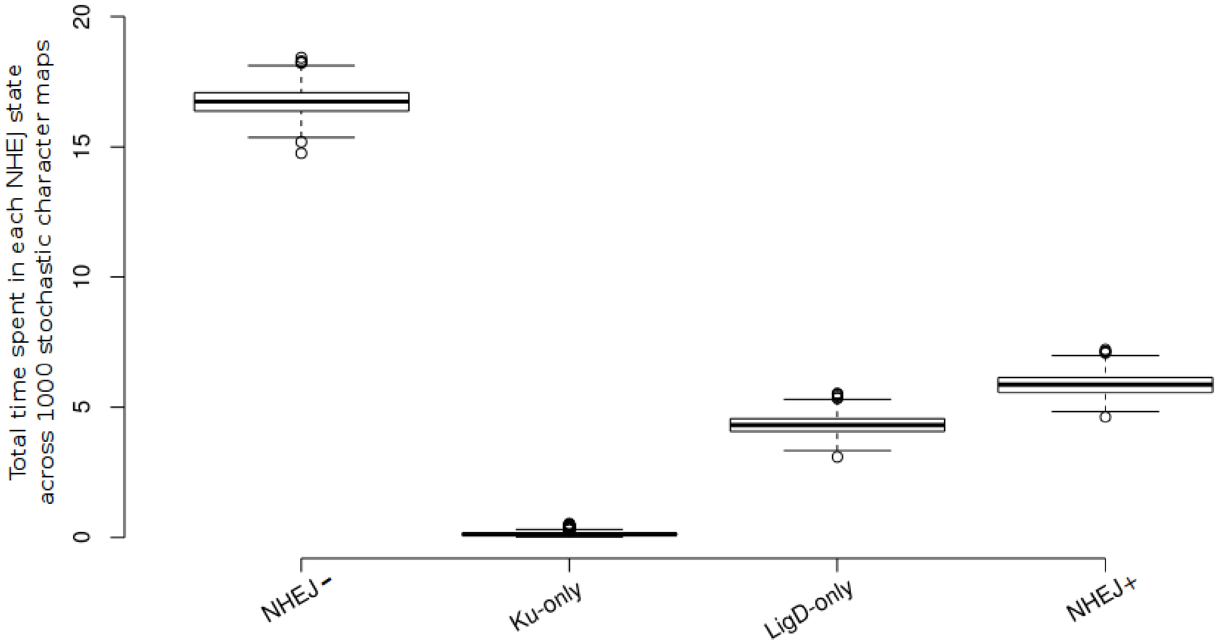


*Figure S1:* Boxplots representing the distribution of total time spent in a given NHEJ state over a phylogenetic tree across 1000 stochastic character maps (Y-axis). X-axis represents the four NHEJ states: *NHEJ-*, *Ku-only*, *LigD-only* and *NHEJ+*.

**FigureS2**

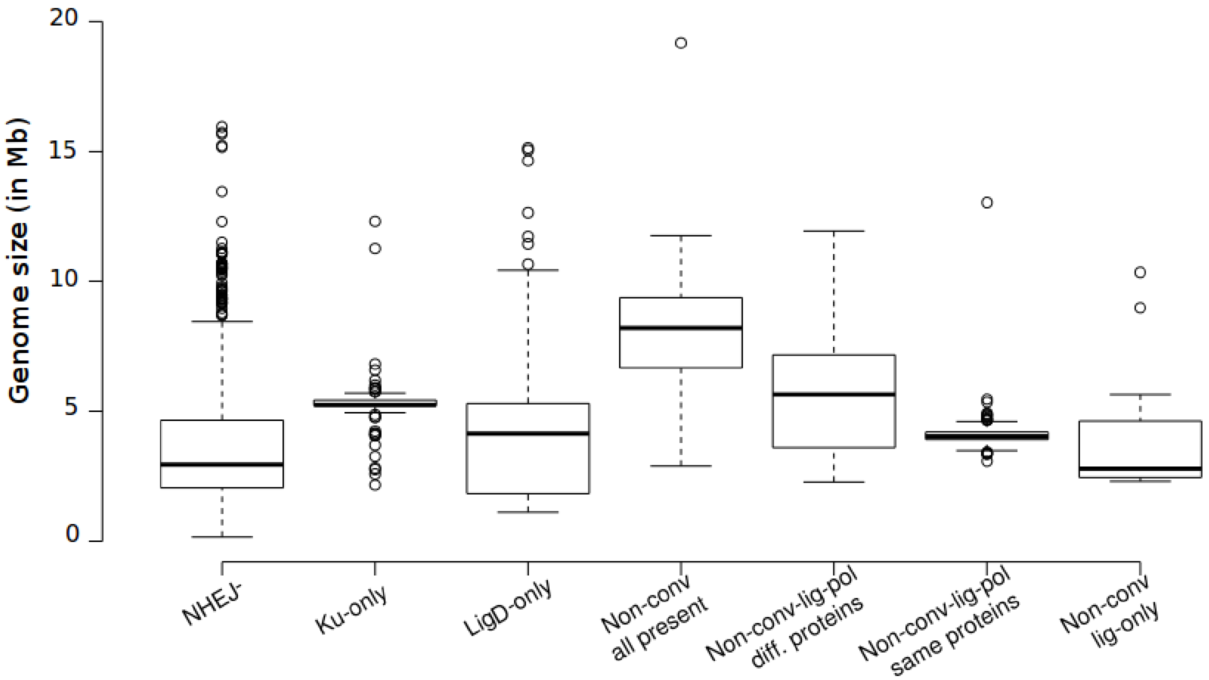


*Figure S2:*Boxplots representing distribution of genome sizes in bacteria with different combinations of NHEJ machinery present: 1) *NHEJ-*, 2) *Ku-only*, 3) *LigD-only*, 4) *Non-conventional NHEJ* (all domains present excluding the conventional LigD), 5) *Non-conventional NHEJ* ( LIG and POL domain present in different proteins and no PE domain), 6) *Non-conventional NHEJ* (LIG and POL domain present in the same protein and no PE domain) and 7) *Non-conventional NHEJ* (LIG domain present and no POL domain)

**Figure S3**

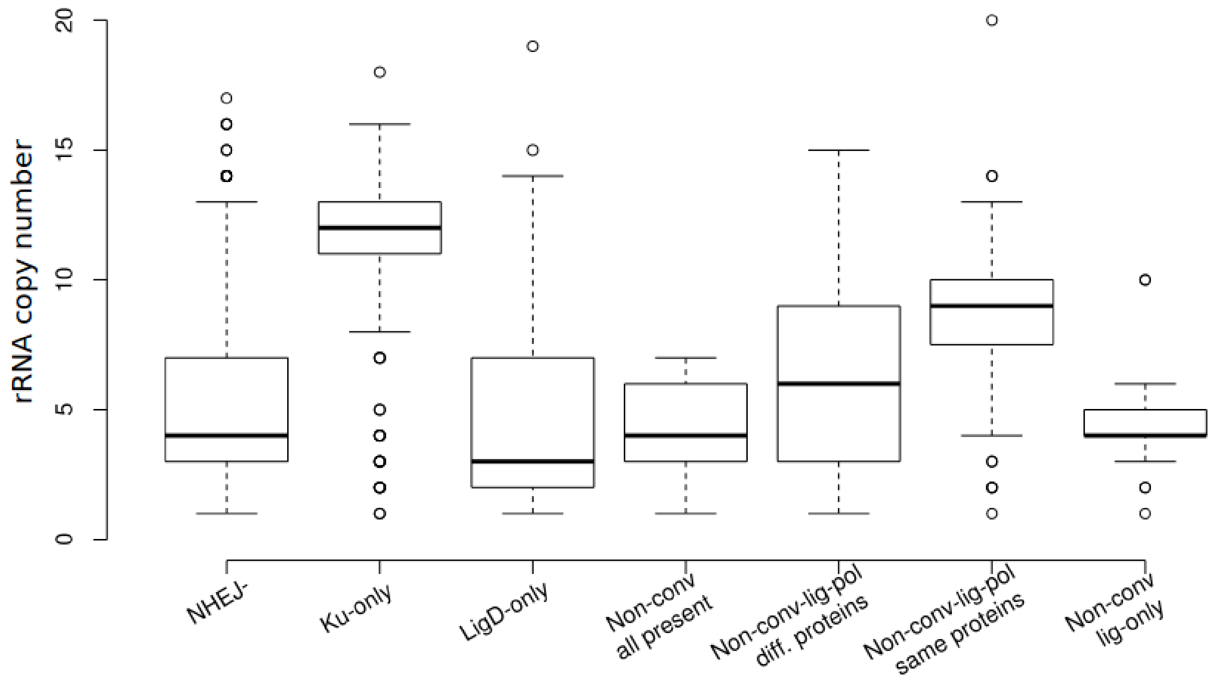


*Figure S3:* Boxplots representing distribution of rRNA copy numbers in bacteria with different combinations of NHEJ machinery present: 1) *NHEJ-*, 2) *Ku-only*, 3) *LigD-only*, 4) *Non-conventional NHEJ* (all domains present excluding the conventional LigD), 5) *Non-conventional NHEJ* ( LIG and POL domain present in different proteins and no PE domain), 6) *Non-conventional NHEJ* (LIG and POL domain present in the same protein and no PE domain) and 7) *Non-conventional NHEJ* (LIG domain present and no POL domain)

**Figure S4**

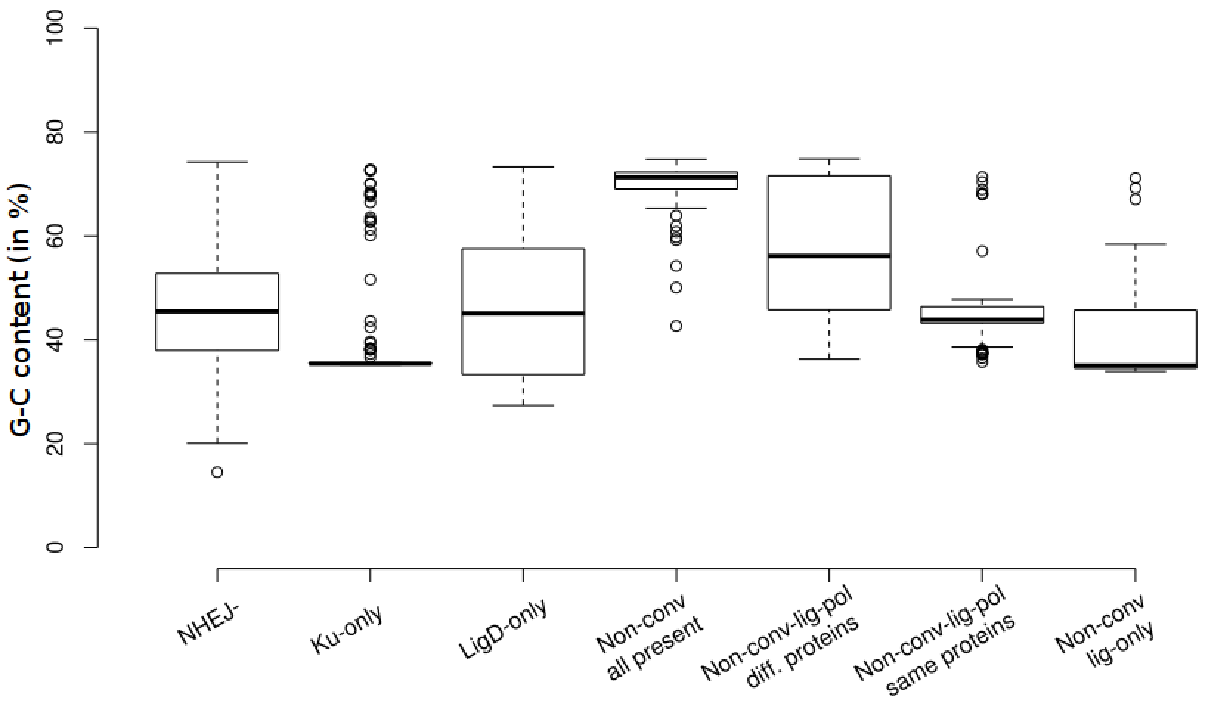


*Figure S4:* Boxplots representing distribution of G-C content in bacteria with different combinations of NHEJ machinery present: 1) *NHEJ-*, 2) *Ku-only*, 3) *LigD-only*, 4) *Non-conventional NHEJ* (all domains present excluding the conventional LigD), 5) *Non-conventional NHEJ* ( LIG and POL domain present in different proteins and no PE domain), 6) *Non-conventional NHEJ* (LIG and POL domain present in the same protein and no PE domain) and 7) *Non-conventional NHEJ* (LIG domain present and no POL domain)

**Figure S5**

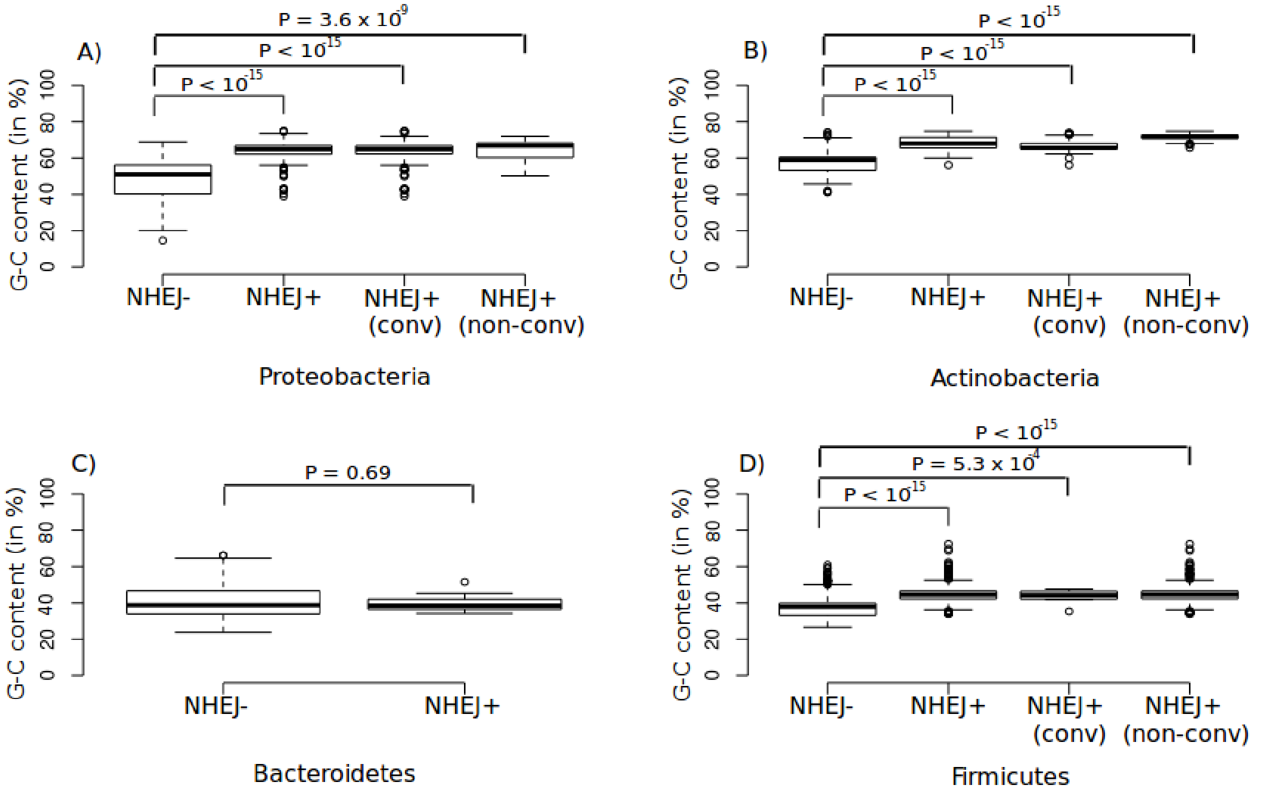


*Figure S5:* Boxplots representing distribution of G-C content in bacteria with different NHEJ states - *NHEJ-* , *conventional NHEJ+* and *non-conventional NHEJ+* across four phyla. G-C content is significantly higher in organisms with NHEJ as compared to those that lack it in three phyla – A) Proteobacteria, B) Actinobacteria and D) Firmicutes. There is no significance difference in G-C content in C) Bacteroidetes.

**Figure S6**

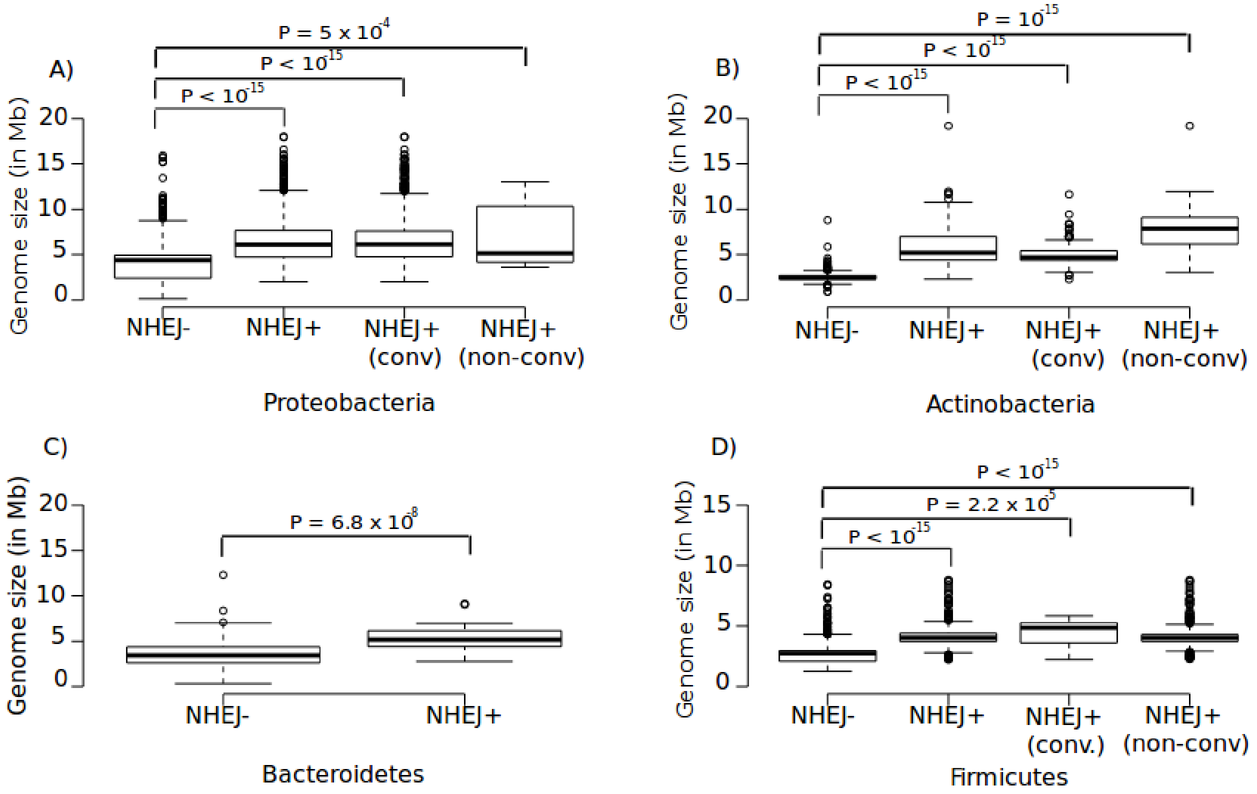


*Figure S6:* Boxplots representing distribution of genome size in bacteria with different NHEJ states - *NHEJ-* , *conventional NHEJ+* and *non-conventional NHEJ+* across four phyla. Genome size is significantly higher in organisms with NHEJ as compared to those that lack it in all four phyla – A) Proteobacteria, B) Actinobacteria, C) Bacteroidetes and D) Firmicutes.

**Figure S7**

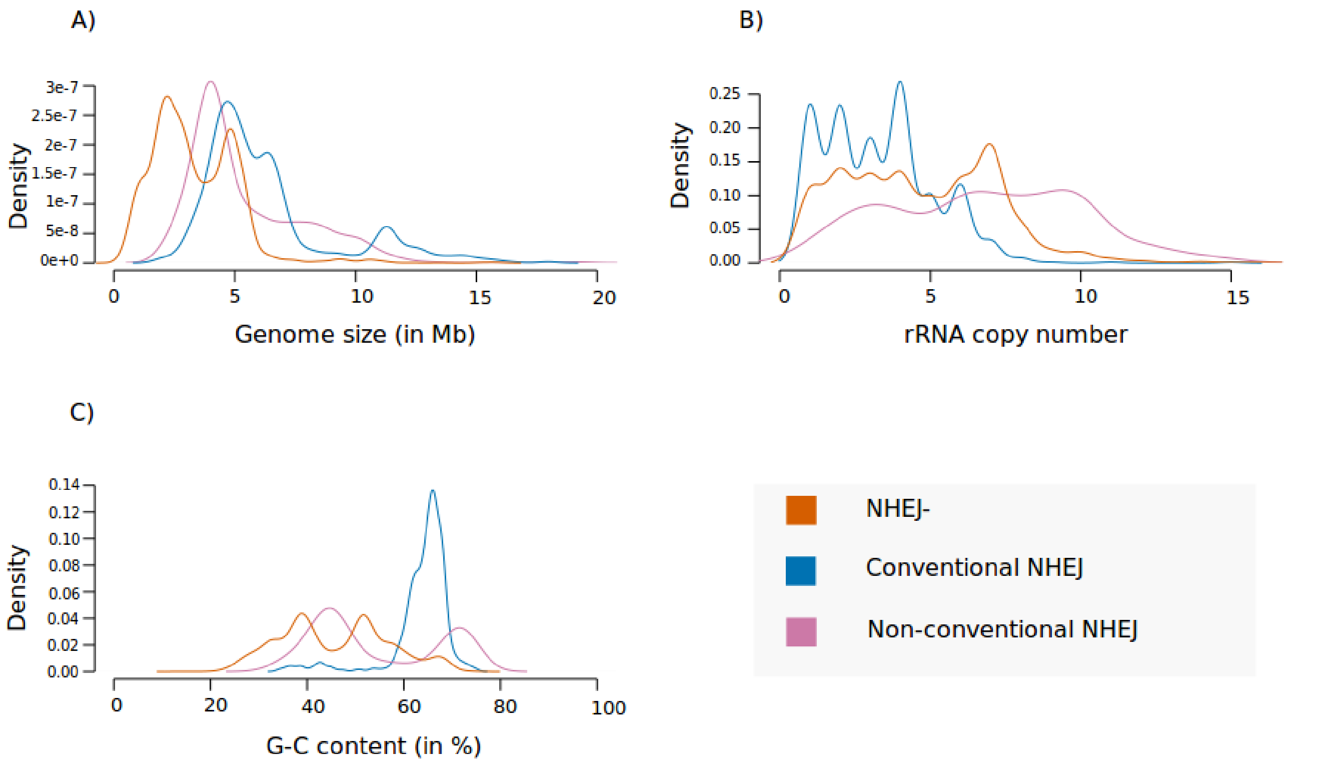


*Figure S7:* Density plots representing the distribution of A) Genome size, B) rRNA copy number and C) G-C content across bacteria with different NHEJ states- *NHEJ-* (orange), *Conventional NHEJ* (blue) and *Non-conventional NHEJ* (pink).

**Figure S8**

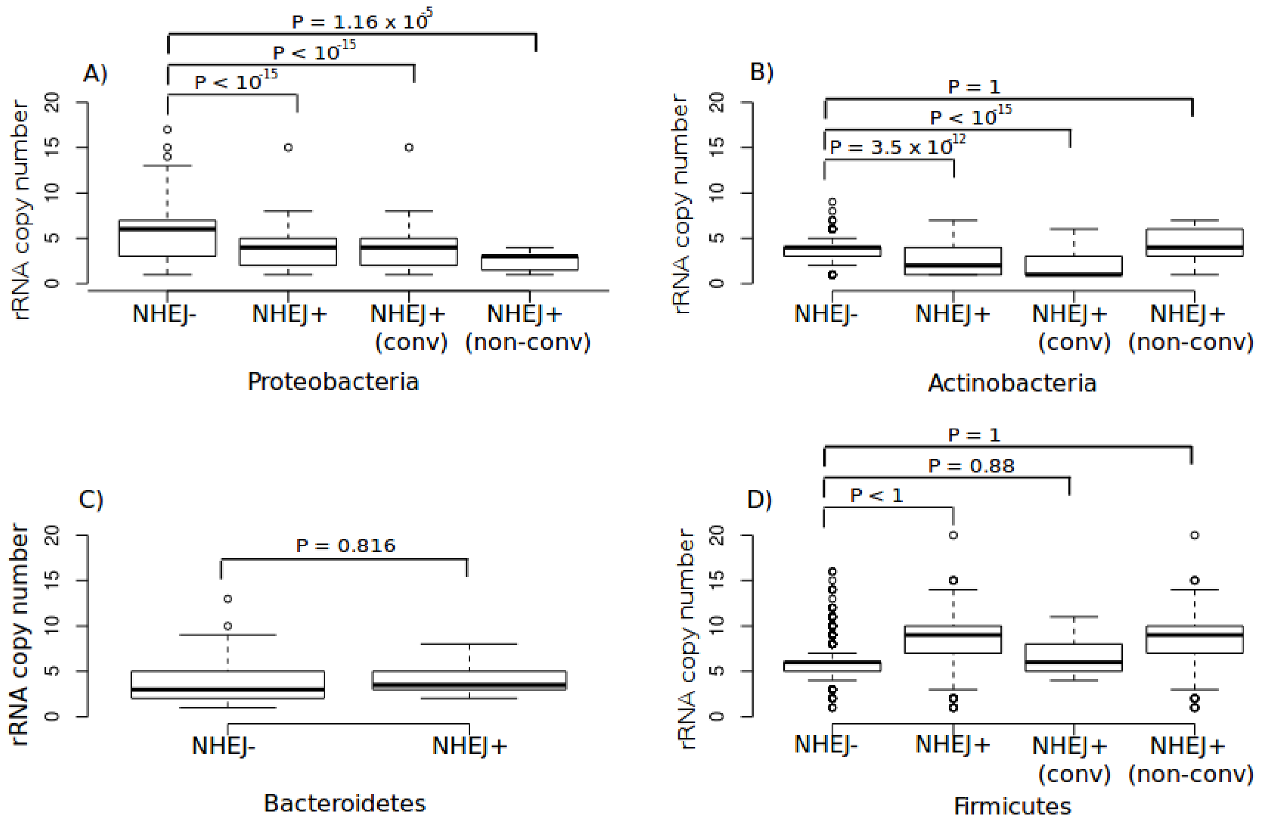


*Figure S8:* Boxplots representing distribution of rRNA copy number in bacteria with different NHEJ states - *NHEJ-* , *conventional NHEJ+* and *non-conventional NHEJ+* across four phyla. rRNA copy number is significantly lower in organisms with NHEJ as compared to those that lack it in two phyla – A) Proteobacteria, B) Actinobacteria. There is no significant difference in rRNA copy number distributions in C) Bacteroidetes. In D) Firmicutes, organisms harboring NHEJ (conventional and non-conventional) tend to have significantly higher growth rate than organisms without NHEJ. However, when *NHEJ +* organisms are segregated into conventional and non-conventional, we find no significant difference in rRNA copy numbers between *NHEJ–* and *conventional NHEJ* Firmicutes.

**Figure S9**

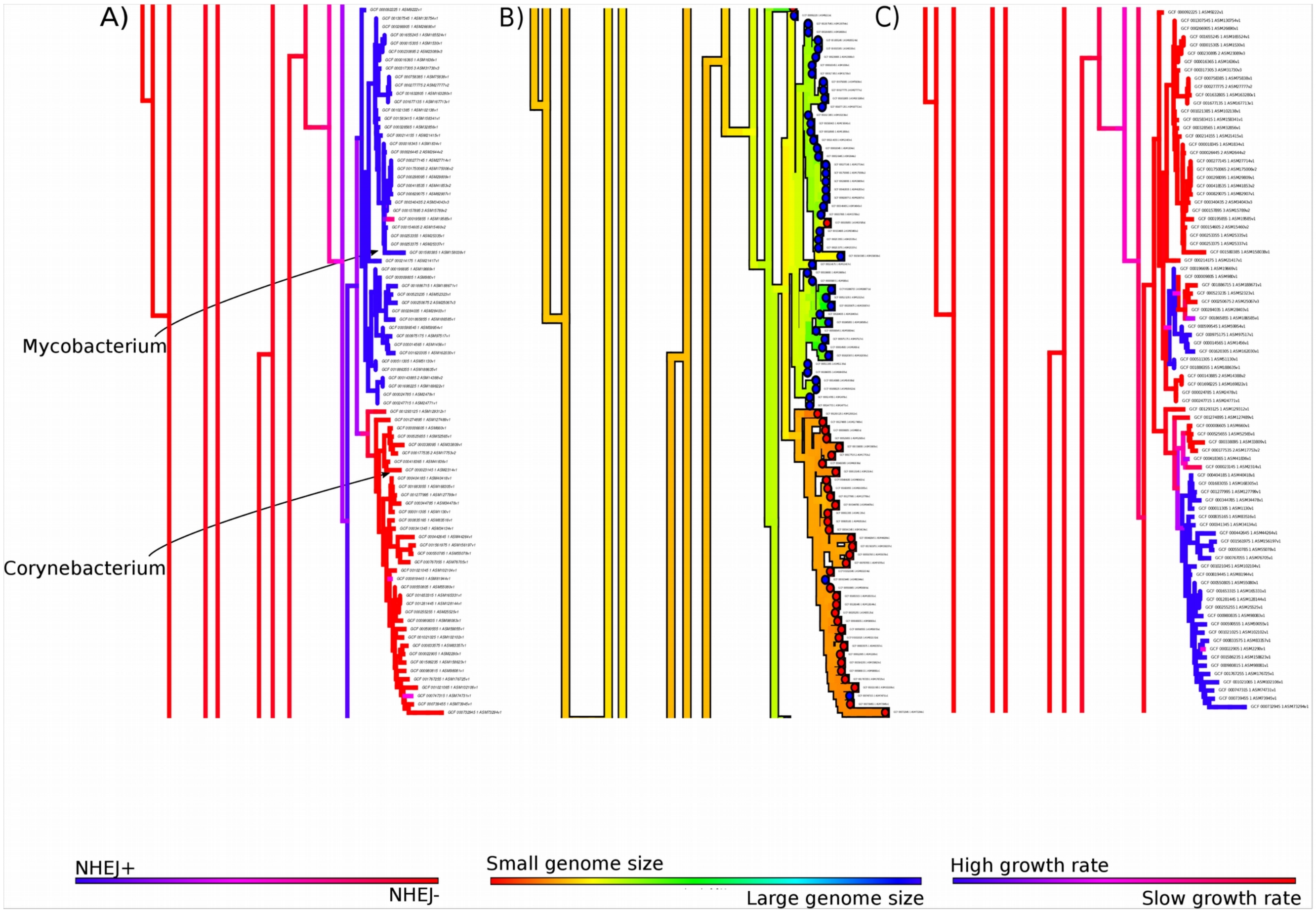


*Figure S9:* A) Density map of phylogenetic ancestral reconstruction of binary trait *conventional NHEJ* presence (blue) and absence (red) represented as posterior probability across 1000 stochastic maps using the All Rates Different (ARD) model, in two Corynebacteriales sub-clades: *Mycobacterium* and *Corynebacterium*. B) Density map of phylogenetic ancestral reconstruction of the continuous trait genome size using the Brownian motion model. The warmer colours represent smaller genome size and colder colours represent larger genome sizes. The tip colours represent conventional NHEJ presence (blue) and absence (red) in the contemporaneous species. C) Density map of phylogenetic ancestral reconstruction of the binary trait growth rate represented as posterior probability across 1000 stochastic maps using All Rates Different (ARD) model.

**Figure S10**

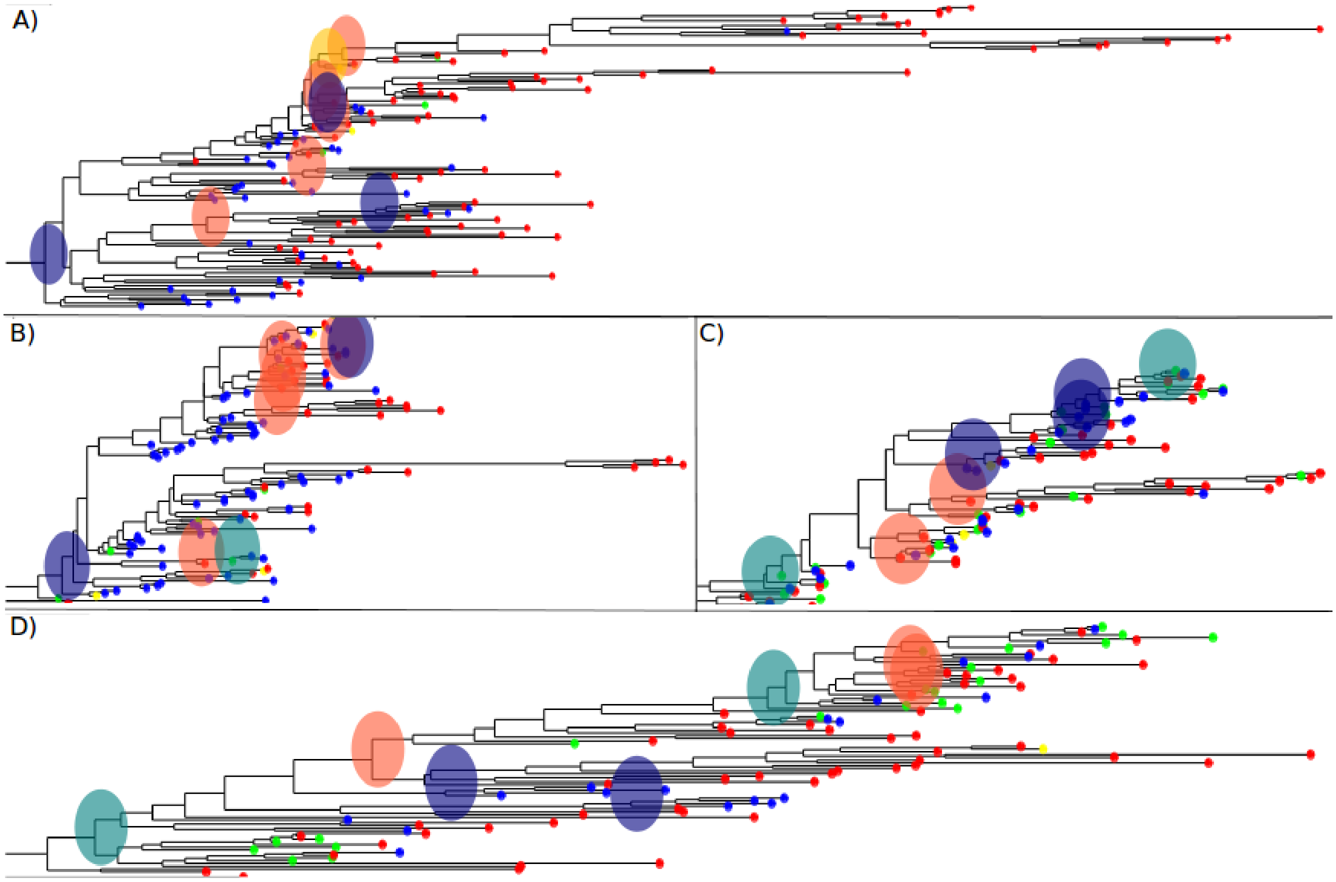


*Figure S10:* A) Direct primary NHEJ gain at the ancestral node of Firmicutes, Fusobacteria and Tenericutes (shallowest phylogenetic depth). B) Direct primary NHEJ gain at the ancestral node of Actinobacteria. C) Sequential primary gain of LigD followed by Ku to yield NHEJ in sub-clade of Proteobacteria. D) Sequential primary gain of LigD in the ancestral node of Bacteroidetes, followed by the complete gain of NHEJ by the acquisition of Ku in different sub-clades. Tip and Node colours- Red: *NHEJ-* , Blue: *NHEJ+* , Green: *LigD only* , Yellow: *Ku only*.
