## Supplementary_table_7 for "Evolutionary and Comparative analysis of bacterial Non-Homologous End Joining Repair"

**Supplementary table 7**
**Contingency tables and Fisher's Exact test results, run on 969 organisms:**

!Conventional NHEJ = Non-conventional NHEJ + No-NHEJ
!Non-conventional NHEJ = Conventional NHEJ + No-NHEJ

| | Proteobacteria | ! Proteobacteria |
|---|---|---|
| Conventional NHEJ | 98 | 58 |
| ! Conventional NHEJ | 368 | 446 |

```
Fisher's Exact Test for Count Data

data:  prot.conventional
p-value = 3.803e-05
alternative hypothesis: true odds ratio is greater than 1
95 percent confidence interval:
 1.501099      Inf
sample estimates:
odds ratio
  2.046295
```

| | Proteobacteria | ! Proteobacteria |
|---|---|---|
| Non-Conventional NHEJ | 15 | 91 |
| ! Non-Conventional NHEJ | 451 | 413 |

```
Fisher's Exact Test for Count Data

data:  prot.nonconventional
p-value = 1
alternative hypothesis: true odds ratio is greater than 1
95 percent confidence interval:
 0.08864445        Inf
sample estimates:
odds ratio
  0.151218
```

| | Actinobacteria | ! Actinobacteria |
|---|---|---|
| Conventional NHEJ | 28 | 128 |
| ! Conventional NHEJ | 716 | 98 |

```
Fisher's Exact Test for Count Data

data:  actino.conventional
p-value = 1
alternative hypothesis: true odds ratio is greater than 1
95 percent confidence interval:
 0.01976974        Inf
sample estimates:
odds ratio
0.03013512
```

| | Actinobacteria | ! Actinobacteria |
|---|---|---|

| | | |
|---|---|---|
| Non-Conventional NHEJ | 44 | 62 |
| ! Non-Conventional NHEJ | 82 | 782 |

```
Fisher's Exact Test for Count Data

data:  actino.nonconventional
p-value = 2.123e-15
alternative hypothesis: true odds ratio is greater than 1
95 percent confidence interval:
 4.511277      Inf
sample estimates:
odds ratio
  6.745681
```

| | Bacteroidetes | ! Bacteroidetes |
|---|---|---|
| Conventional NHEJ | 19 | 137 |
| ! Conventional NHEJ | 74 | 740 |

```
Fisher's Exact Test for Count Data

data:  bacteroidetes.conventional
p-value = 0.1468
alternative hypothesis: true odds ratio is greater than 1
95 percent confidence interval:
 0.8406495       Inf
sample estimates:
odds ratio
   1.38635
```

| | Bacteroidetes | ! Bacteroidetes |
|---|---|---|
| Non-Conventional NHEJ | 0 | 106 |
| ! Non-Conventional NHEJ | 93 | 771 |

```
Fisher's Exact Test for Count Data

data:  bacteroidetes.nonconventional
p-value = 1
alternative hypothesis: true odds ratio is greater than 1
95 percent confidence interval:
   0 Inf
sample estimates:
odds ratio
          0
```

| | Firmicutes | ! Firmicutes |
|---|---|---|
| Conventional NHEJ | 5 | 151 |
| ! Conventional NHEJ | 122 | 692 |

```
Fisher's Exact Test for Count Data

data:  firmicutes.conventional
p-value = 1
```

```
alternative hypothesis: true odds ratio is greater than 1
95 percent confidence interval:
 0.07174517        Inf
sample estimates:
odds ratio
 0.1880178
```

| | Firmicutes | ! Firmicutes |
|---|---|---|
| Non-Conventional NHEJ | 42 | 64 |
| ! Non-Conventional NHEJ | 85 | 779 |

```
Fisher's Exact Test for Count Data

data:  firmicutes.nonconventional
p-value = 1.23e-13
alternative hypothesis: true odds ratio is greater than 1
95 percent confidence interval:
 4.00504     Inf
sample estimates:
odds ratio
  5.997159
```

| | Acidobacteria | ! Acidobacteria |
|---|---|---|
| Conventional NHEJ | 3 | 153 |
| ! Conventional NHEJ | 3 | 811 |

```
Fisher's Exact Test for Count Data

data:  acidobacteria.conventional
p-value = 0.05622
alternative hypothesis: true odds ratio is greater than 1
95 percent confidence interval:
 0.947979      Inf
sample estimates:
odds ratio
  5.286889
```

| | Acidobacteria | ! Acidobacteria |
|---|---|---|
| Non-Conventional NHEJ | 2 | 104 |
| ! Non-Conventional NHEJ | 4 | 860 |

```
Fisher's Exact Test for Count Data

data:  acidobacteria.nonconventional
p-value = 0.1325
alternative hypothesis: true odds ratio is greater than 1
95 percent confidence interval:
 0.5472948       Inf
sample estimates:
odds ratio
  4.125139
```

| | Chlamydiae | ! Chlamydiae |
|---|---|---|
| Conventional NHEJ | 1 | 155 |
| ! Conventional NHEJ | 5 | 809 |

```
Fisher's Exact Test for Count Data

data:  chlamydiae.conventional
p-value = 0.6518
alternative hypothesis: true odds ratio is greater than 1
95 percent confidence interval:
 0.04453014        Inf
sample estimates:
odds ratio
  1.043831
```

| | Chlamydiae | ! Chlamydiae |
|---|---|---|
| Non-Conventional NHEJ | 1 | 105 |
| ! Non-Conventional NHEJ | 5 | 859 |

```
Fisher's Exact Test for Count Data

data:  chlamydiae.nonconventional
p-value = 0.5015
alternative hypothesis: true odds ratio is greater than 1
95 percent confidence interval:
 0.06959043        Inf
sample estimates:
odds ratio
  1.635181
```

| | Verrucomicrobia | ! Verrucomicrobia |
|---|---|---|
| Conventional NHEJ | 0 | 156 |
| ! Conventional NHEJ | 6 | 808 |

```
Fisher's Exact Test for Count Data

data:  verrucomicrobia.conventional
p-value = 1
alternative hypothesis: true odds ratio is greater than 1
95 percent confidence interval:
   0 Inf
sample estimates:
odds ratio
         0
```

| | Verrucomicrobia | ! Verrucomicrobia |
|---|---|---|
| Non-Conventional NHEJ | 1 | 105 |
| ! Non-Conventional NHEJ | 5 | 859 |

```
Fisher's Exact Test for Count Data

data:  verrucomicrobia.nonconventional
p-value = 0.5015
alternative hypothesis: true odds ratio is greater than 1
95 percent confidence interval:
 0.06959043        Inf
sample estimates:
odds ratio
  1.635181
```

-----------------------------------------------------------------------------------------------------------------------

**List of phyla with all *No-NHEJ* state organisms only:**

1) Tenericutes
2) Thermotogae
3) Synergistetes
4) Fusobacteria
5) Chlorobi
6) Deferribacteres
7) Elusimicrobia
8) Ignavibacteriae
9) Fibrobacteres
10) Coprothermobacterota
11) Chrysiogenetes
12) Calditrichaeota
13) Kirtimatiellaeota
14) Candidatus Cloacimonetes

-----------------------------------------------------------------------------------------------------------------------

**List of phyla with *incomplete NHEJ* or *No-NHEJ* state organisms only:**

1) Caldiserica
2) Dictyoglomi
3) Gemmatimonadetes
4) Thermobaculum
5) Deinococcus-Thermus
6) Planctomycetes
7) Aquificae
8) Chloroflexi
9) Cyanobacteria
10) Spirochaetes
