## Supplementary_table_8_9 for "Evolutionary and Comparative analysis of bacterial Non-Homologous End Joining Repair"

| | Sub-phyla name | Type of gain | Type of loss | Organism names |
|---|---|---|---|---|
| | Flavobacteriia | Sequential secondary gain (NHEJ- to LigD to NHEJ+) | - | Maribacter sp. 1_2014MBL_MicDiv; Cellulophaga baltica NN016038; Flavobacterium johnsoniae UW101; Aequorivita sublithincola DSM 14238; Zunongwangia profunda SM-A87; Gramella sp. LPB0144 |
| | | Secondary direct gain (NHEJ- to NHEJ+) | - | Chryseobacterium sp. IHB B17019; Flavobacteriaceae bacterium 3519-10 |
| | Common ancestor of Flavobacteriia and Bacteroidia | - | Major Secondary loss of LigD to NHEJ- | - |
| | Sphingobacteria | Primary sequential gain (LigD to NHEJ+) | - | Mucilaginibacter sp. PAMC 26640 plasmid unnamed; Pedobacter saltans DSM 12145; Sphingobacterium sp. ML3W; Solitalea canadensis DSM 3403 |
| | Chitinophagia | Primary NHEJ sequential gain (LigD to NHEJ+) | - | Niabella ginsenosidivorans strain BS26; Niastella koreensis GR20−10; Flavisolibacter sp. LCS9; Chitinophaga pinensis DSM 2588 |
| | Cytophagia | Primary NHEJ sequential gain (LigD to NHEJ+) | - | Cytophaga hutchinsonii ATCC 33406; Dyadobacter fermentans DSM 18053; Cyclobacterium amurskyense strain KCTC 12363 |
| | Common ancestor of Flavobacteriia, Bacteroidia, Sphingobacteria, Chitinophagia and a subclade of Cytophagia | Major Primary LigD gain | - | - |

| | | | | |
|---|---|---|---|---|
| | Thermoleophilia | Minor Primary direct gain (NHEJ- to NHEJ+) | - | Conexibacter woesei DSM 14684 |
| | Coriobacteriia | Minor Primary direct gain (NHEJ- to NHEJ+) | - | Eggerthella lenta DSM 2243 |
| | Acidimicrobiia | Minor Primary direct gain (NHEJ- to NHEJ+) | - | Ilumatobacter coccineus YM16−304 |
| | Common ancestor of Frankiales, Geodermatophilales, Streptosporangiales, Streptomycetales, Catenulisporales, Glycomycetales, Pseudonocardiales, Propionibacteriales, Corynebacteriales, Micromonosporales, Bifidobacteriales, Micrococcales, Kineosporiales, Nakamurellales, Actinomycetales | Major Primary direct gain | - | |
| | Frankiales | Primary direct gain | | Frankia sp. EuI1c; Frankia sp. EAN1pec |
| | Geodermatophilales | Primary direct gain | | Blastococcus saxobsidens DD2; Geodermatophilus obscurus DSM 43160; Modestobacter marinus str. BC501 |
| | Actinobacteria incertae sedis | Primary direct gain | | Thermobispora bispora DSM 43833 |
| | Streptosporangiales | Primary direct gain | | Thermomonospora curvata DSM 43183; Streptosporangium roseum DSM 43021 |
| | Streptosporangiales | - | Direct secondary loss | Thermobifida fusca; Nocardiopsis alba ATCC BAA−2165 |
| | Catenulisporales | Primary direct gain | | Catenulispora acidiphila DSM 44928 |
| | Streptomycetales | - | Direct secondary loss | Kitasatospora setae KM−6054 DNA |
| | Streptomycetales | Primary direct gain | - | Streptomyces lydicus strain 103; |

| | | | | |
|---|---|---|---|---|
| | | | | Streptomyces sampsonii strain KJ40 |
| | Pseudonocardiales | Primary direct gain | - | Actinoalloteichus hymeniacidonis strain HPA177(T) (=DSM 45092(T)); Pseudonocardia dioxanivorans CB1190; Kibdelosporangium phytohabitans strain KLBMP1111; Saccharothrix espanaensis DSM; Alloactinosynnema sp. L−07; Actinosynnema mirum DSM 43827; Lentzea guizhouensis strain DHS C013; Saccharopolyspora erythraea NRRL2338; Saccharomonospora viridis DSM 43017; Amycolatopsis japonica strain MG417−CF17; |
| | Glycomycetales | Primary direct gain | - | Stackebrandtia nassauensis DSM 44728; |
| | Corynebacteriales | Primary direct gain | - | Rhodococcus fascians D188; Nocardia seriolae strain EM150506; Corynebacterium humireducens NBRC 106098 = DSM 45392; Mycobacterium tuberculosis 49−02; Mycobacterium goodii strain 7B; Amycolicicoccus subflavus DQS3−9A1; Gordonia sp. QH−11; Gordonia sp. KTR9; Tsukamurella paurometabola DSM 20162; |
| | Corynebacteriales | - | Direct secondary loss | Brevibacterium flavum ZL−1; Corynebacteriales bacterium 1698; Segniliparus rotundus DSM 44985; Corynebacterium pseudotuberculosis strain 36; |

| | | | | |
|---|---|---|---|---|
| | | | | Mycobacterium leprae TN |
| | Propionibacteriales | Direct primary gain | - | Microlunatus phosphovorus NM−1 DNA;<br>Nocardioides dokdonensis FR1436;<br>Pimelobacter simplex strain VKM Ac−2033D;<br>Aeromicrobium erythreum strain AR18;<br>Kribbella flavida DSM 17836 |
| | Propionibacteriales | - | Direct secondary loss | Cutibacterium avidum strain DPC 6544;<br>Propionibacterium acnes C1 |
| | Bifidobacteriales | - | Direct secondary loss | Scardovia inopinata JCM 12537 DNA;<br>Parascardovia denticolens DSM 10105 = JCM 12538 DNA;<br>Gardnerella vaginalis ATCC 14018 = JCM 11026 DNA;<br>Bifidobacterium breve 12L |
| | Micromonosporales | Direct primary gain | - | Salinispora arenicola CNS−205;<br>Salinispora tropica CNB−440;<br>Micromonospora sp. L5;<br>Verrucosispora maris AB−18−032;<br>Actinoplanes friuliensis DSM 7358 |
| | Nakamurellales | Direct primary gain | - | Nakamurella multipartita DSM 44233 |
| | Kineosporiales | Direct primary gain | - | Kineococcus radiotolerans SRS30216 plasmid pKRAD02 |
| | Micrococcales | Direct primary gain | - | Xylanimonas cellulosilytica DSM 15894 ;<br>Isoptericola dokdonensis DS−3;<br>Beutenbergia cavernae DSM 12333;<br>Serinicoccus sp. JLT9;<br>Kytococcus sedentarius DSM 20547;<br>Dermacoccus |

| | | | | |
|---|---|---|---|---|
| | | | | nishinomiyaensis strain M25; Luteipulveratus mongoliensis strain MN07−A0370; Arsenicicoccus sp. oral taxon 190; Intrasporangium calvum DSM 43043; Janibacter terrae strain \| F\|001 |
| | Micrococcales | - | Direct secondary loss | Jonesia denitrificans DSM 20603; Tropheryma whipplei TW08/27 |
| | Actinomycetales | - | Direct secondary loss | Trueperella pyogenes TP8; Arcanobacterium haemolyticum DSM 20595; Actinobaculum schaalii strain CCUG 27420; Actinomyces sp. Marseille−P2985 strain Marseille−P2985T ; Mobiluncus curtisii ATCC 43063; |
| | Micrococcales | Direct primary gain | - | Brevibacterium linens strain SMQ−1335; Sanguibacter keddieii DSM 10542; Cellvibrio gilvus ATCC 13127; Cellulomonas fimi ATCC 484; Brachybacterium faecium DSM 4810; Sinomonas atrocyanea strain KCTC 3377; Arthrobacter aurescens TC1 plasmid TC2; Rathayibacter tritici strain NCPPB 1953; Agromyces aureus strain AR33; Microbacterium sp. 1.5R; Microbacterium sp. T11; Cryobacterium arcticum strain PAMC 27867; Curtobacterium sp. BH−2−1−1; Frondihabitans sp. SR6 plasmid 3; Clavibacter |

| | | | | |
|---|---|---|---|---|
| | | | | michiganensis subsp. insidiosus strain R1−1; Leifsonia xyli strain SE134 |
| | Micrococcales | - | Direct secondary loss | Devriesea agamarum genome assembly S2_rcS3_S1_rcS3; Rothia mucilaginosa D\|−18 DNA; Kocuria rhizophila DC2201; Dermabacter vaginalis strain AD1−86; Neomicrococcus aestuarii strain B18; Arthrobacter sp. A3; Renibacterium salmoninarum ATCC 33209; Micrococcus luteus NCTC 2665; Microbacterium sp. No. 7; Rathayibacter toxicus strain WAC3373; Microterricola viridarii strain ERGS5:02 |
| | Fimbriimonadia | Direct primary gain | - | Fimbriimonas ginsengisoli Gsoil 348 |
| | Clostridia | Direct primary gain | - | Thermaerobacter marianensis DSM 12885; Candidatus Desulforudis audaxviator MP104C; |
| | Thermodesulfobacteriales | Direct primary gain | - | Thermodesulfatator indicus DSM 15286 |
| | Clostridia | Direct primary gain | - | Thermosediminibacter oceani DSM 16646; Thermacetogenium phaeum DSM 12270; Carboxydothermus hydrogenoformans Z−2901; Syntrophothermus lipocalidus DSM 12680; Syntrophomonas wolfei subsp. wolfei str. Goettingen G311; Moorella thermoacetica strain DSM 103132; Natranaerobius thermophilus |

| | | | | |
|---|---|---|---|---|
| | | | | JW/NM−WN−LF; Thermoanaerobacter sp. 513; Halothermothrix orenii H 168; Limnochorda pilosa DNA; Symbiobacterium thermophilum IAM 14863 DNA; Clostridium cellulosi genome assembly DG5; Clostridium stercorarium subsp. stercorarium DSM 8532; Clostridium stercorarium subsp. leptospartum DSM 9219; Thermoanaerobacterium thermosaccharolyticum M0795; Desulfotomaculum reducens MI−1; Thermincola sp. JR; Desulfitobacterium dehalogenans ATCC 51507; Desulfitobacterium metallireducens DSM 15288; Syntrophobotulus glycolicus DSM 8271; Heliobacterium modesticaldum Ice1 strain Ice1; Desulfosporosinus meridiei DSM 13257; Dehalobacter restrictus DSM 9455; Pelotomaculum thermopropionicum SI DNA |
| | Clostridia | - | Direct Secondary loss | Tepidanaerobacter acetatoxydans Re1; Caldicelulosiruptor becscii DSM 6725; Caldicellulosiruptor saccharolyticus DSM 8903; Ethanoligenens harbinense \|UAN−3; Ruminococcus bicirculans; Halobacteroides halobius DSM 5150; Acetohalobium |

| | | | | |
|---|---|---|---|---|
| | | | | arabaticum DSM 5501; Halanaerobium praevalens DSM 2228; Flavonifractor plautii strain \|L31; Intestinimonas butyriciproducens strain AF211; Oscillibacter valericigenes Sjm18−20; Mahella australiensis 50−1 BON; Ruminiclostridium thermocellum DSM 2360; Clostridium clariflavum DSM 19732; Clostridiales genomosp. BVAB3 str. UPII9−5; Eubacterium limosum strain SA11; Acetobacterium woodii DSM 1030; Desulfotomaculum acetoxidans DSM 771 |
| | Common ancestor of Tissierellia and a sub-clade of Clostridia | - | Direct Secondary loss | |
| | Clostridia | - | Direct Secondary loss | Clostridium propionicum DSM 1682; Peptoclostridium difficile strain 08ACD0030; Filifactor alocis ATCC 35896; Geosporobacter ferrireducens strain IRF9; Alkaliphilus metalliredigens Q\|MF; Lachnoclostridium sp. \| L32; Butyrivibrio proteoclasticus B316; Roseburia hominis A2−183 |
| | Clostridia | Direct Secondary gain | - | Blautia sp. \|L58; Clostridium saccharolyticum WM1; Clostridium phytofermentans ISDg |

| | | | | |
|---|---|---|---|---|
| | Tissierellia | - | Direct Secondary loss | Levyella sp. Marseille−P3170 strain Marseille−P3170T ; Murdochiella sp. Marseille−P2341 strain Marseille−P2341T ; Parvimonas micra strain KCOM 1535; Finegoldia magna ATCC 29328; Peptoniphilus sp. ING2−D1G; Anaerococcus prevotii DSM 20548 |
| | Negativicutes | - | Direct Secondary loss | Acidaminococcus fermentans DSM 20731; Selenomonas sputigena ATCC 35185; Dialister pneumosintes strain F0677; Megasphaera elsdenii 14−14; Veillonella parvula DSM 2008 |
| | | Direct Secondary gain | - | Pelosinus fermentans JBW45 |
| | Bacilli | Direct Primary gain | - | Kyrpidia tusciae DSM 2912; Alicyclobacillus acidocaldarius subsp. acidocaldarius Tc−4−1; Thermobacillus composti KWC4; Bacillus cellulosilyticus DSM 2522; Paenibacillus sp. LPB0068; Paenibacillus polymyxa SC2; Halobacillus halophilus DSM 2266; Terribacillus aidingensis strain MP602; Lentibacillus amyloliquefaciens strain LAM0015; Oceanobacillus iheyensis HTE831; Virgibacillus sp. 6R; Bacillus pseudofirmus OF4; Lysinibacillus sphaericus strain 2362; Solibacillus silvestris strain DSM 12223; Rummeliibacillus stabekisii strain PP9; |

| | | | | |
|---|---|---|---|---|
| | | - | Direct Secondary loss | Geobacillus stearothermophilus 10; Paenibacillus sp. BD3526; Salimicrobium jeotgali strain MJ3; Amphibacillus xylanus NBRC 15112; Planococcus rifietoensis strain M8; Anoxybacillus sp. B7M1; Exiguobacterium antarcticum B7; Sporosarcina psychrophila strain DSM 6497; Aneurinibacillus sp. H2; Kurthia sp. 11Kri321; Jeotgalibacillus sp. D5; Leuconostoc carnosum JB16; Oenococcus oeni PSU−1; Weissella koreensis KACC 15510; Pediococcus damnosus strain TMW 2.1534; Aerococcus christensenii strain CCUG28831; Lactobacillus salivarius strain JCM 1046; Marinilactibacillus sp. 15R; Carnobacterium sp. WN1359; Streptococcus constellatus subsp. pharyngis C818; Lactococcus garvieae Lg2 DNA; Enterococcus faecium strain 64/3; Melissococcus plutonius S1; Vagococcus teuberi strain DSM21459T; Tetragenococcus halophilus NBRC 12172 DNA; Listeria monocytogenes strain ATCC 19117; Macrococcus caseolyticus JCSC5402; Salinicoccus halodurans |

| | | | | |
|---|---|---|---|---|
| | | | | strain H3B36; Bacillus cereus subsp. cytotoxis NVH 391−98; Gemella sp. oral taxon 928; Staphylococcus aureus strain CA15 |
| | | Direct Secondary gain | - | Brevibacillus laterosporus LMG 15441 |
| | Erysipelotrichia | Direct Secondary gain | - | Erysipelotrichaceae bacterium I46 |
| | | - | Direct Secondary loss | Turicibacter sp. H121; Erysipelothrix rhusiopathiae str. Fujisawa DNA; Faecalibaculum rodentium strain Alo17; |
| | Fusobacteriales | - | Direct Secondary loss | Ilyobacter polytropus DSM 2926; Fusobacterium nucleatum subsp. animalis strain KCOM 1279v; Sebaldella termitidis ATCC 33386; Leptotrichia sp. oral taxon 847; Streptobacillus moniliformis DSM 12112; Sneathia sp. Sn35; |
| | Mollicutes | - | Direct Secondary loss | Strawberry lethal yellows phytoplasma (CPA) str. NZSb11; Aster yellows witches'−broom phytoplasma A\|WB; Maize bushy stunt phytoplasma strain M3; Onion yellows phytoplasma O\|−M DNA; Acholeplasma palmae; Acholeplasma oculi; Acholeplasma oculi strain 19L; Mollicutes bacterium HR1; Mycoplasma mycoides subsp. mycoides strain Ben468; Mesoplasma florum L1; Spiroplasma culicicola |

| | | | | |
|---|---|---|---|---|
| | | | | AES−1; Ureaplasma urealyticum serovar 10 str. ATCC 33699 |
| | Nitrospirales | Direct primary gain | - | Nitrospira moscoviensis strain NSP M−1 |
| | Solibacteres | Direct primary gain | - | Solibacter usitatus Ellin6076 |
| | Acidobacteriales | Direct primary gain | - | Candidatus Koribacter versatilis Ellin345; Terriglobus saanensis SP1PR4; Acidobacterium capsulatum ATCC 51196; Granulicella mallensis MP5ACT 8; |
| | Parachlamydiales | Direct primary gain | - | Protochlamydia naegleriophila genome assembly PNK1; Parachlamydia acanthamoebae UV−7 |
| | Opitutae | Direct primary gain | - | Opitutus terrae PB90−1 |
| | Deltaproteobacteria | Direct primary gain | - | Desulfovibrio africanus str. Walvis Bay; Desulfomonile tiedjei DSM 6799; Geobacter sp. M21; Geobacter uraniireducens Rf4; Myxococcus fulvus 124B02; Myxococcus stipitatus DSM 14675; Archangium gephyra strain DSM 2261; Vulgatibacter incomptus strain DSM 27710; Anaeromyxobacter dehalogenans 2CP−1; Sorangium cellulosum So0157−2; Sorangium cellulosum 'So ce 56'; Chondromyces crocatus strain Cm c5; Sandaracinus amylolyticus strain DSM 53668; Haliangium ochraceum DSM 14365 |
| | Alphaproteobacteria | Direct primary gain | - | Gluconacetobacter diazotrophicus PAl 5; |

| | | | | |
|---|---|---|---|---|
| | | | | Tistrella mobilis KA081020-065; Porphyrobacter neustonensis strain DSM 9434; Citromicrobium sp. JL477; Erythrobacter litoralis strain DSM 8509; Altererythrobacter atlanticus strain 26DY36; Croceicoccus naphthovorans strain PQ-2; Sphingopyxis terrae NBRC 15098 strain 203-1; Blastomonas sp. RAC04; Blastomonas sp. RAC04; Novosphingobium aromaticivorans DSM 12444; Sphingomonas melonis; Sphingomonas sanxanigenens DSM 19645; Sphingobium sp. SYK−6; Caulobacteraceae bacterium OTSz_A_272; Caulobacter sp. K31; Phenylobacterium zucineum HLK1; Brevundimonas sp. GW460−12−10−14−LB 2; Asticcacaulis excentricus CB 48; Bradyrhizobium icense strain LMTR 13; Rhodopseudomonas palustris BisB5; Oligotropha carboxidovorans OM5; Nitrobacter winogradskyi Nb−255; Methylocella silvestris BL2; Beijerinckia indica subsp. indica ATCC 9039; Methylocystis sp. SC2; Starkeya novella DSM 506 anthobacter |

| | | | | |
|---|---|---|---|---|
| | | | | autotrophicus Py2; Azorhizobium caulinodans ORS 571; Rhodoplanes sp. Z2−\| C6860; Chelatococcus daeguensis strain TAD1; Bosea vaviloviae strain Vaf18; Mesorhizobium loti MAFF303099; Chelativorans sp. BNC1; Martelella endophytica strain \| C6887; Aminobacter aminovorans strain KCTC 2477; Hoeflea sp. IMCC20628 ; Neorhizobium galegae chromid pHAMBI540a; Agrobacterium tumefaciens strain S33; Ensifer adhaerens strain Casida A; Sinorhizobium americanum CCGM7; Ochrobactrum anthropi strain OAB; Filomicrobium sp. ; Hyphomicrobium denitrificans ATCC 51888; Rhodomicrobium vannielii ATCC 17100; Methyloceanibacter caenitepidi; Devosia sp. H5989; Pelagibacterium halotolerans B2; Rhizobium phaseoli strain R650; Shinella sp. HZN7; Parvibaculum lavamentivorans DS−1; |
| | | - | Direct secondary loss | Alpha proteobacterium HIMB59; Novosphingobium pentaromativorans US6−1; Sphingopyxis sp. LPB0140; Zymomonas mobilis subsp. pomaceae ATCC 29192; Maricaulis maris MCS10; Micavibrio aeruginosavorus EPB; Magnetospirillum sp. |

| | | | | |
|---|---|---|---|---|
| | | | | M−1; Rhodospirillum rubrum ATCC 11170; Haematospirillum jordaniae strain H5569; Thalassospira xiamenensis M−5 = DSM 17429; Liberibacter crescens BT-1; Brucella abortus strain 63 75; Bartonella tribocorum; Magnetospira sp. QH−2 |
| | | Sequential secondary gain | - | Defluviimonas alba strain cai42; Rhodobacter sphaeroides ATCC 17025; Sulfitobacter sp. AM1-D1; Celeribacter indicus strain P73 |
| | Gammaproteobacteria | Direct primary gain | - | Luteibacter rhizovicinus strain LJ96T; Lysobacter capsici strain 55; Lysobacter antibioticus strain ATCC 29479; Dokdonella koreensis DS−123; Dyella jiangningensis strain SBZ 3−12; Pseudoxanthomonas spadix BD−a59; Stenotrophomonas maltophilia D457; Xanthomonas gardneri strain ICMP 7383; |
| | | - | Direct secondary loss | Xylella fastidiosa 9a5c plasmid pXF51; Xanthomonas albilineans str. GPE PC73; |
| | Betaproteobacteria | Direct primary gain | - | Nitrosospira briensis C−128; Methylovorus glucosetrophus SIP3-4; Massilia sp. WG5 plasmid unnamed 2 |
| | | Sequential primary gain | - | Janthinobacterium sp. 1_2014MBL_MicDiv; Pandoraea apista strain DSM 16535; Pandoraea sputorum strain DSM 21091;Collimonas |

| | | | | |
|---|---|---|---|---|
| | | | | fungivorans Ter331; Herbaspirillum seropedicae SmR1; Paraburkholderia caribensis strain Bcrs1W; Burkholderia pseudomallei Pasteur 52237; Burkholderia pseudomallei strain Burk178-Type2; Azoarcus sp. KH32C plasmid pAZKH DNA; Achromobacter xylosoxidans genome assembly NCTC10807; Bordetella flabilis strain AU10664; Bordetella holmesii ATCC 51541; Advenella kashmirensis WT001; Polyangium brachysporum strain DSM 7029; Rubrivivax gelatinosus IL144 DNA; Delftia sp. HK171; Mitsuaria sp. 7; Roseateles depolymerans strain KCTC 42856; Acidovorax ebreus TPS\|; Acidovorax citrulli AAC00-1; Ramlibacter tataouinensis TTB310; Variovorax paradoxus EPS; Polaromonas sp. JS666; Hydrogenophaga sp. PBC; |
| | | Direct tertiary gain | - | Cupriavidus basilensis strain 4G11; Ralstonia mannitolilytica strain SN82F48; Thiobacillus denitrificans ATCC 25259; |
| | | - | Direct secondary loss | Basilea psittacipulmonis DSM 24701; Alcaligenes faecalis strain ZD02; Castellaniella; defragrans 65Phen; Bordetella pertussis strain E476; Pusillimonas sp. T7−7; Taylorella equigenitalis ATCC 35865; |

| | | | | |
|---|---|---|---|---|
| | Gammaproteobacteria | Direct primary gain | - | Legionella hackeliae genome assembly LHA; Tatlockia micdadei genome assembly LMI; Coxiella burnetii CbuG_Q212; Pseudomonas syringae pv. tomato str. DC3000; Pseudomonas stutzeri DSM 4166; Halomonas chromatireducens strain AGD 8−3 |
| | | - | Direct secondary loss | Mutant Legionella pneumophila subsp. pneumophila str.; Hextuple_3a Legionella pneumophila str. Corby; |
| | | Direct secondary gain | - | Pseudomonas aeruginosa strain PA_D16 |

Note: This table doesn't contain *NHEJ*- to *LigD/Ku* events that do not culminate to NHEJ+ eventually and *LigD/Ku* to *NHEJ*- events. Other organisms not listed in the table do not code for NHEJ by the virtue of the eubacterial ancestor.

**Supplementary table 9**

| Variable | Test statistics | K value | λ value | $logL_0$ | logL | P-value |
|---|---|---|---|---|---|---|
| Genome size | Pagel's λ | - | 0.91 | 95.07 | 426.25 | $4.56x10^{-146}$ |
| Genome size | Blomberg's K | 0.0144 | - | - | - | $1x10^{-3}$ |
| rRNA copy number | Pagel's λ | - | 0.899 | -179.85 | 183.493 | $4.68x10^{-160}$ |
| rRNA copy number | Blomberg's K | 0.00368 | - | - | - | $1x10^{-3}$ |

**Table depicting the strength of phylogenetic signal for genome size and rRNA copy number across a phylogenetic tree of 969 bacteria using two test statistics – Pagel's λ and Blomberg's K - employed for continuous character traits.** Both the measures suggest that phylogenetic conservatism is significantly greater than random expectations for both the genome characteristics, albeit there is a discordance between the strength of phylogenetic signal for both the genome characteristics measured by Pagel's λ and Blomberg's K.
